# Chromosome-level genome assembly of the primitive loach goby, *Rhyacichthys aspro*, reveals mechanisms underlying Gobioidei diversification

**DOI:** 10.1101/2024.04.02.587698

**Authors:** Tzi-Yuan Wang, Hao-Jun Lu, Yu-Wei Wu, Te-Yu Liao, Shih-Pin Huang, Feng-Yu Wang, Chau-Ti Ting, Shu-Miaw Chaw, Hurng-Yi Wang

**Affiliations:** Biodiversity Research Center, Academia Sinica, Nankang, Taipei, Taiwan; Department of Life Science, National Taiwan University, Taipei, Taiwan; Graduate Institute of Clinical Medicine, College of Medicine, National Taiwan University, Taipei, Taiwan; Graduate Institute of Biomedical Informatics, College of Medical Science and Technology, Taipei Medical University, Taipei, Taiwan; Department of Oceanography, National Sun Yat-sen University, Kaohsiung, Taiwan; Taiwan Ocean Research Institute, National Applied Research Laboratories, Kaohsiung, Taiwan; Institute of Ecology and Evolutionary Biology, National Taiwan University, Taipei 10617, Taiwan; Graduate Institute of Medical Genomics and Proteomics, College of Medicine, National Taiwan University, Taipei 10002, Taiwan; Department of Entomology, National Taiwan University, Taipei 10617, Taiwan

**Keywords:** Loach goby, Transposons, Repeat elements, Adaptation, Genome evolution, Diversification

## Abstract

The percomorph fish clade Gobioidei are a suborder that comprises over 2,200 species distributed in nearly all aquatic habitats. To understand the genetics underlying their diversification, we sequenced and annotated the genome of the loach goby, *Rhyacichthys aspro*, the basal most group, and compared it with nine additional Gobioidei species. Within Gobioidei, the loach goby possesses the smallest genome at 607 Mb, and a rise in species diversity from basal to derived lineages is mirrored by enlarged genomes and a higher presence of repeat elements (REs), particularly DNA transposons. These transposons are enriched in coding and regulatory regions and their copy number increase is strongly correlated with mutation rate, suggesting that DNA repair after transposon excision/insertion leads to nearby mutations. Consequently, the proliferation of DNA transposons might be the crucial driver of Gobioidei diversification and adaptability. The loach goby genome also points to mechanisms of ecological adaptation. It contains relatively few genes for lateral line development but an over representation of synaptic function genes, with genes putatively under selection linked to synapse organization and calcium signaling, suggesting a sensory system distinct from other Gobioidei species. We also see an overabundance of genes involved in neurocranium development and renal function, adaptations likely connected to its flat morphology suited for strong currents and an amphidromous life cycle. Comparative analyses with hill-stream loaches and the European eel reveal convergent adaptations in body shape and saltwater balance. These findings shed light on the loach goby’s survival mechanisms and the broader evolutionary trends within Gobioidei.

## 1 INTRODUCTION

The order Gobiiformes represent a remarkable array of diversity within vertebrates, encompassing over 2,200 species across 321 genera, ranking it among the largest vertebrate orders (Kuang et al., 2018; Nelson, Grande, & Wilson, 2016). These fish occupy all aquatic environments on earth (Brandl, Goatley, Bellwood, & Tornabene, 2018; Brandl et al., 2019). Most species are marine, while around 10% are found in freshwater habitats (Sidlauskas & Chakrabarty, 2010). Characterized by their extraordinary variety of forms, ecological roles, and behaviors, Gobiiformes have become prominent model organisms in comparative research, shedding light on the evolutionary mechanisms driving species diversification (Agorreta et al., 2013; de Brito et al., 2022). The order Gobiiformes includes nine families. The Rhyacichthyidae, Odontobutidae, Milyeringidae, Eleotridae, Butidae, Thalasseleotrididae, Oxudercidae, and Gobiidae belong to the suborder Gobioidei, while the Trichonotidae, the only family in suborder Trichonotoidei, are the sister group of all species in the suborder Gobioidei (Nelson et al., 2016; Christine E. Thacker et al., 2015; C. E. Thacker et al., 2023) (Figure 1a). Within Gobioidei, Oxudercidae and Gobiidae are the most derived families, comprising over 1,957 species and 275 genera (Nelson et al., 2016).

**Figure 1.**
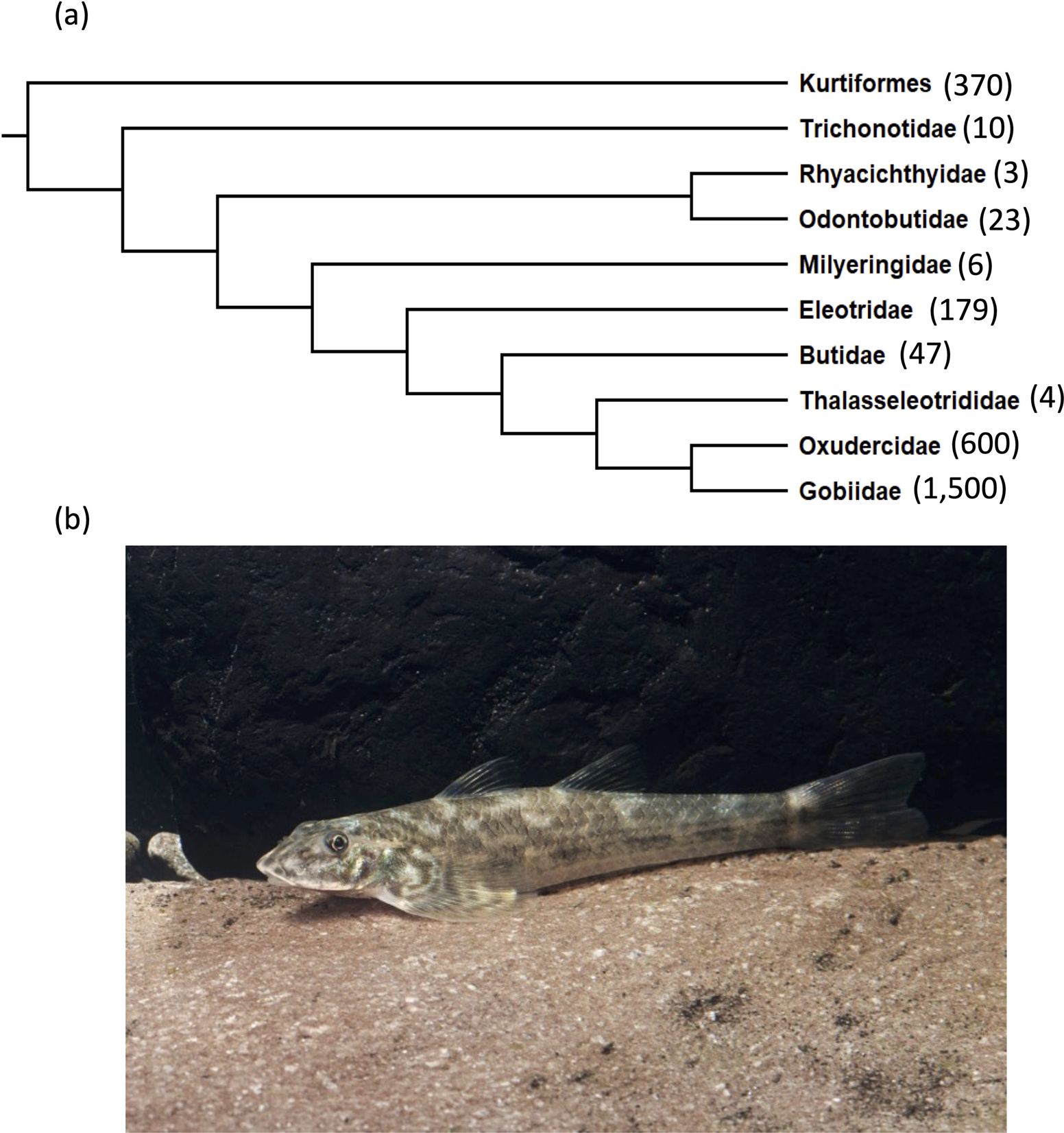
Relationships and diversity in Gobiiformes. (a) Phylogeny of Gobiiformes with Kurtiformes as an outgroup. Numbers in parenthesis are putative species number. The Milyeringidae and Thalasseleotrididae were formerly classified as part of the family Eleotridae but researchers had noted that these genera were atypical members of the Eleotridae. (b) Photo of *Rhyacichthys aspro* taken by Dr. Te-Yu Liao.

Gobioidei are a highly diverse suborder with adaptations to a wide range of environments. Therefore, understanding the genetic basis of these adaptations will shed light on the evolutionary processes that underpin species diversification and the mechanisms that enable such a vast array of species to flourish in both marine and freshwater ecosystems. Access to genome sequences of an organism offers unparalleled insights into its genetic blueprint, supporting the elucidation of its evolutionary history, biological functions, and molecular mechanisms. Moreover, genomic data provides a foundation for comparative studies, helping identify unique adaptations, ecological interactions, and evolutionary innovations. While many gobioid genome assemblies have been generated recently (Adrian-Kalchhauser et al., 2020; Cai et al., 2021; Y. Hu et al., 2021; Q. Liu et al., 2021; Y. Yang et al., 2022; You et al., 2014), a reference genome for the earliest divergent family of Gobioidei, the Rhyacichthyidae, is yet to be assembled. Analysis and annotation of a representative Rhyacichthyidae genome will enable comparison to other recently diverged Gobioidei species and should offer invaluable perspectives on the evolution and diversification of this suborder.

Within Gobioidei, Rhyacichthyidae exhibit several comparatively primitive characteristics, such as three epurals, six branchiostegal rays, and unfused pelvic fins, that differentiate it from derived relatives (Hoese & Gill, 1993; Miller, 1973). Perhaps the most distinct feature of Rhyacichthyidae is its sensory system. Lateral-line systems of the vast majority of gobioid fishes are reduced to cephalic canals, accompanied by a remarkable proliferation of petite superficial neuromasts (located on ‘sensory papillae’). This unique sensory system may indicate derived gobies have specialized neuron and sensory development. The Rhyacichthyidae, however, do not possess this specialized feature. Instead, they have a well-developed lateral line that runs along the head and body (Akihito et al., 2000; Hoese & Gill, 1993; Miller, 1973).

The Rhyacichthyidae comprise only two genera and three species including *Rhyacichthys aspro* (Cuvier and Valenciennes 1837) (Figure 1b), commonly known as the loach goby, that can be found throughout the Indo-Australasian archipelago, China, Taiwan, and Japan (Froese & Pauly, 2023). Adult loach gobies inhabit freshwater streams of typically rocky or gravelly areas with fast-flowing water. Morphologically, the species is characterized by an elongated, cylindrical body with a notably flattened head and snout. Loach gobies inhabit benthic environments, primarily consuming diatoms and algae. A previous study based on otoliths suggests that the species is amphidromous (Tabouret et al., 2014) migrating between freshwater and marine environments to complete its life cycle.

To elucidate the genomic basis of species diversification and evolution in Gobioidei, we sequenced, assembled, and annotated a loach goby, *Rhyacichthys aspro*, genome from northern Taiwan. We then performed a comparative genomic analysis using nine additional available genomes representing four Gobioidei families (Table 1). Our results show that expansion of repeat elements (REs), especially DNA transposon, is associated with increased mutation rate and species diversification of Gobioidei, suggesting that DNA transposon activity is a key factor in the extensive diversity and adaptability of Gobioidei. Additionally, we identified unique gene sets related to synaptic development, neurocranium morphogenesis, and osmoregulation. These findings illuminate the organism’s adaptations to sensory demands, turbulent currents, and the amphidromous nature of its habitats.

**Table 1.**
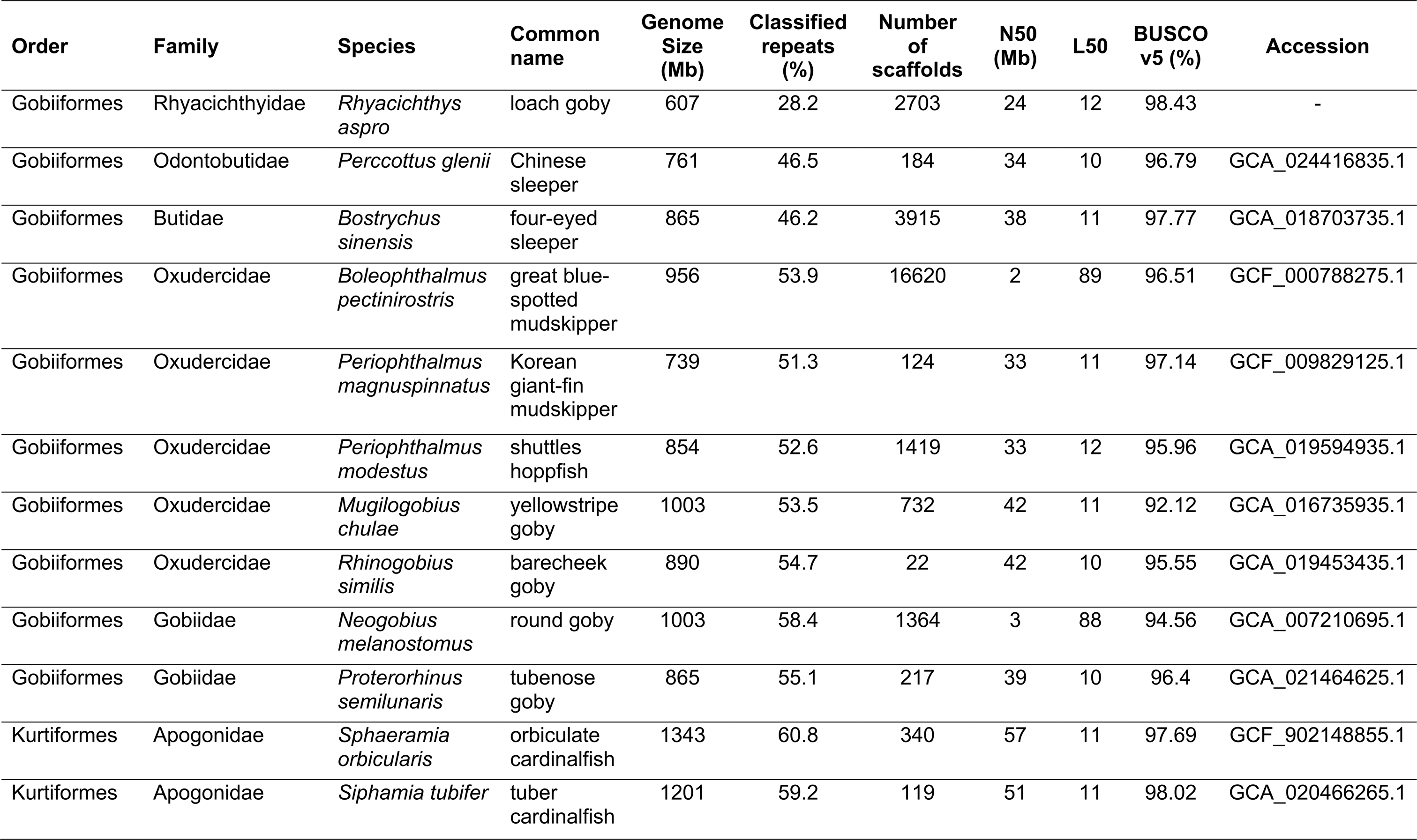
Genome statistics of the selected species.

## 2 MATERIALS and METHODS

### 2.1 Sample collection

An adult loach goby, *Rhyacichthys aspro*, was collected from Masu creek (Masu1) (25°10’16.4“N 121°40’47.7“E), New Taipei, Taiwan. The head was dissected from fish body and preserved in RNAlater™ Stabilization Solution (Cat: AM7021, Invitrogen, Thermo Fisher Scientific Inc.) at -80°C freezer. The other tissues were preserved in 100% ethanol at -80°C freezer.

To investigate nucleotide diversity and population size of the loach goby in Taiwan, four additional individuals were collected and sequenced. Two individuals (Gangkou1 and Gangkou2) were collected from Gangkou creek, Pingtung, Taiwan, one (JhonggongM) from Jhonggong creek, Miaoli, and sample (JhonggongP) from Jhonggong creek, Pingtung (Table S1). All experiments in this study were performed in accordance with guidelines of the animal ethics committee and were approved by the Institutional Animal Care and Use Committee (IACUC 20-12-1593), Academia Sinica.

### 2.2 DNA and RNA sequencing

30-50 mg of muscle tissue was lysed with proteinase K and Buffer G2 (with RNAse A) of Genomic DNA Buffer Set (Cat: 19060, Qiagen) and the genomic DNA was extracted via Genomic-tip 20/G (Cat: 10223, Qiagen), following the manufacturer’s instructions.

Genomic DNA (5∼8 μg) for Nanopore sequencing was end-repaired and ligated via a KAPA Hyper Prep Kit (Cat#KR0961, Kapa Biosystems, Wilmington, MA, USA), following the manufacturer’s instructions. The genomic DNA library was premixed with the LB and SQB buffer in the Nanopore Ligation Sequencing Kit (SQK-LSK109, Oxford Nanopore Technologies, UK) loaded in a flow cell (R9.4.1; FLO-MIN106), and sequenced by MinION devices for 24-72 h.

Genomic DNA was sheared using a Covaris 2 machine, and then quantified using a Qubit 2.0 Fluorometer before Illumina sequencing. A short-read sequencing library was also constructed from 200ng of the sheared gDNA using the TruSeq Nano DNA High Throughput Library Prep Kit (Cat. No.20015965). It was then sequenced using an Illumina X-ten through the GENOMICS BIOSCIENCE & TECHNOLOGY CO., LTD service based in Taipei, Taiwan.

Five to 30 mg of eye, olfactory lobe, gill, liver, muscle, and brain tissues were dissected and homogenized with 600 µl Trizol solution for RNA sequencing. RNA was extracted from the homogenized Trizol solution using RNeasy® Plus Mini Kit (Cat: 74134, Qiagen), and quantified using the Agilent 2100 Bioanalyzer. A short-read sequencing library was constructed from approximately 1.5μg of the quantified RNA using the TruSeq Stranded mRNA Library Prep Kit (Cat. No. 20020595). It was then sequenced on the Illumina NovaSeq 6000 through the services of GENOMICS.

### 2.3 Draft genome assembly

Nanopore long reads were assembled using canu v1.8 (Koren et al., 2017) with “-nanopore-raw” parameter. The resulting assembly was error-corrected using: 1) racon v1.4.3 (Vaser, Sovic, Nagarajan, & Sikic, 2017) on canu-corrected Nanopore long reads four times; 2) medaka v0.11.2 (https://github.com/nanoporetech/medaka) on canu-corrected Nanopore long reads once; and 3) pilon v1.23 (Walker et al., 2014) on trimmed Illumina short reads (using Trimmomatic v0.39 (Bolger, Lohse, & Usadel, 2014) with the following parameters: illuminaclip:truseq3-pe.fa:2:30:10 leading:20 trailing:20 slidingwindow:4:15 minlen:36.

### 2.4 Chicago and HiRise scaffolding

Both Chicago libraries (Putnam et al., 2016) and Dovetail HiC libraries were prepared as described previously (Lieberman-Aiden et al., 2009). The *de novo* draft assembly, Chicago library reads, and Dovetail HiC library reads were used as input data for HiRise, a software pipeline designed specifically for using proximity ligation data to scaffold genome assemblies (Putnam et al., 2016). An iterative analysis was conducted. First, Chicago library sequences were aligned to the draft input assembly using a modified SNAP read mapper (http://snap.cs.berkeley.edu). The separations of Chicago read pairs mapped within draft scaffolds were analyzed by HiRise to produce a likelihood model for genomic distance between read pairs, and the model was used to identify and break putative misjoins, to score prospective joins, and make joins above a threshold. After aligning and scaffolding Chicago data, Dovetail HiC library sequences were aligned and scaffolded following the same method.

### 2.5 Genome quality analysis and genome masking

The genome quality was calculated using BUSCO v5 (Manni, Berkeley, Seppey, Simão, & Zdobnov, 2021) score in actinopterygii_odb10. BlobTools v1.1 (Laetsch & Blaxter, 2017) was used to check for assembly contamination. The genome size was estimated using Jellyfish (Marcais & Kingsford, 2012). To characterize repeat elements (REs), a consensus library was built by merging the output from a *de novo* library from RepeatModeler2.0.2 (Flynn et al., 2020), an LTR library from LTR_finder (Xu & Wang, 2007) and LTR retriever (Ou & Jiang, 2017), the TE_library from RepBase20181026 (Bao, Kojima, & Kohany, 2015) and Dfam3.5 (Storer, Hubley, Rosen, Wheeler, & Smit, 2021), and the *P. magnuspinnatus* TE library from FishTEDB (Shao, Wang, Xu, & Peng, 2018). REs were masked using RepeatMasker 4.1.2 (Nishimura, 2000). A python script “TE_divergence_landscape.py” (https://github.com/PlantDr430/CSU_scripts/blob/master/TE_divergence_landscape.py) on Github was used to calculate Kimura divergence and create a repeat landscape. Because different databases have been utilized to identify REs of different species in the literature, we reanalyzed the REs of the genomes listed in Table 1 to ensure they are compared using the latest database. Computational Analysis of gene Family Evolution (CAFÉ) 5 (De Bie, Cristianini, Demuth, & Hahn, 2006) was used to calculate the expansion and reduction of the TEs based on copy number within each species and the branch lengths in the phylogeny.

### 2.6 Loach goby genome annotation

BRAKER 2.1.6 (Hoff, Lomsadze, Borodovsky, & Stanke, 2019) was used for genome annotation. It predicts genes based on RNA or homologous protein sequences. For RNA-based prediction, transcriptomes from gill, brain, eye, liver, nose, and muscle were mapped to the assembly using hisat2 v2.2.1 (Kim, Paggi, Park, Bennett, & Salzberg, 2019) to create a bam file. BRAKER was run with the genome, RNA bam file, and options “--softmasking” and “--UTR=on”.

For homology protein based prediction, gene sets of 12 Percomorphaceae species, including *P. magnuspinnatus* (GCF_009829125.1), *Boleophthalmus pectinirostris* (GCF_000788275.1), *Sphaeramia orbicularis* (GCF_902148855.1), *Betta splendens* (GCF_900634795.3), *Scophthalmus maximus* (GCF_013347765.1), *Oryzias latipes* (GCF_002234675.1), *Nothobranchius furzeri* (GCF_001465895.1), *Gasterosteus aculeatus* (GCF_016920845.1), *Larimichthys crocea* (GCF_000972845.2), *Syngnathus acus* (GCF_901709675.1), *Thalassophryne amazonica* (GCF_902500255.1) and *Takifugu rubripes* (GCF_901000725.2) were download from NCBI. An orthology database was created based on genome and protein gene sets using the ProHint BRAKER extension. BRAKER was run with the genome sequence, protein orthology database, and the option “--softmasking”. The TSEBRA BRAKER extension was used to combine RNA and proteinbased annotations. The sequences were then blasted against the NCBI database with an E-value cut off of 1.0E^-5^ and any sequences without BLAST hits were filtered out.

Functional annotation, including NCBI BLAST and InterProScan analyses (Jones et al., 2014), was performed using Blast2GO (Conesa et al., 2005). Gene Ontology (GO) annotations were generated with EggNOG-mapper2.1.0 (Cantalapiedra, Hernández-Plaza, Letunic, Bork, & Huerta-Cepas, 2021), employing the EggNOG5.0.2 database (Huerta-Cepas et al., 2019). BLAST searches were carried out against the Actinopterygii taxonomic group in the NCBI nr database. InterProScan queried multiple databases, including CDD (Marchler-Bauer et al., 2010), HAMAP (Lima et al., 2009), PANTHER (Mi et al., 2005), Pfam (Bateman et al., 2004), PIR (Barker et al., 2000), PRINTS (Attwood et al., 2012), ProSite (Hulo et al., 2006), TIGRFAM (Haft et al., 2012), Gene 3D (Lees et al., 2012), SFLD (Akiva et al., 2014), and SuperFamily (Lima et al., 2009). Subsequently, KAAS (Moriya, Itoh, Okuda, Yoshizawa, & Kanehisa, 2007) was used to search KEGG Orthology identifiers (KO numbers) in selected species using the bidirectional best hit (BBH) method and mapped them to the Kyoto Encyclopedia of Genes and Genomes (KEGG) pathways (Kanehisa & Goto, 2000). Comparative gene sets in the KEGG database were selected from *Homo sapiens*, *Mus musculus*, and all 38 fish species. Finally, PfamScan (Mistry, Bateman, & Finn, 2007) was utilized to search for protein domains within the predicted genes using the Pfam database (Bateman et al., 2004).

Synteny relationships between the loach goby and other selected species (*P. magnuspinnatus*, *Periophthalmus modestus* (GCA_019594935.1), and *S. orbicularis*) with chromosome-level genome assemblies were inferred using DAGchainer (Haas, Delcher, Wortman, & Salzberg, 2004) bundled by Synima (Farrer, 2017). Circos 0.69-9 (Krzywinski et al., 2009) were used to visualize chromosome synteny.

### 2.7 Evolutionary rate analysis

Orthofinder2.52 (Emms & Kelly, 2019) was used to identify orthogroups. Single-copy orthogroups were selected and aligned using MACSEv2.05 (Ranwez, Harispe, Delsuc, & Douzery, 2011). iqtree1.6.12 (Nguyen, Schmidt, Von Haeseler, & Minh, 2015) was then used to construct the phylogenetic tree. dN/dS ratios in single copy orthogroups were calculated using codeml in PAML4.9 (Yang et al., 1997) with the free ratio model. The dN tree, dS tree, and dN/dS tree were created from the PAML results. The pairwise dN and dS values among single-copy genes and the pairwise distance among 12S and 16S rDNAs were calculated using MEGA11.0.13 (Tamura, Stecher, & Kumar, 2021). Gene Ontology (GO) term enrichment analysis was conducted for genes that show signs of positive selection in the lineage leading to the loach goby using Fisher’s extract test with FDR < 0.05.

### 2.8 Population genetic tests and population size estimation

To analyze the nucleotide diversity and estimate the population size of our loach goby population, Illumina short reads of Masu1, Gangkou1, Gangkou2, JhonggongM, and JhonggongP were mapped back to the reference genome using BWA0.7.17 (Li, 2013). VCF files were then generated using samtools1.12 and bcftools1.12-26. The nucleotide diversity pi and Tajima’s D statistics were calculated using vcftools with a vcf file merge from five samples using bcftools.

To identify selective sweeps, Tajima’s D values were calculated using vcf.-kit for 5-kb sliding windows in 1-kp steps. Genes with a 1 kb flanking region containing at least one window with negative D was considered as positively selected.

For MSMC (Schiffels & Wang, 2020) analysis, sites with base quality lower than 20 and mapping quality lower than 20 or mappability rate lower than one were ignored. Read depths lower than one-third or higher than twice the average were likewise discarded. Diploid fastq was created using vcfutils.pl from filtered vcf files containing data from each sample. Population sizes were then estimated using the resulting fastq. -p (The patterns of parameters) was set at “1*2+50*1+1*2+1*3”. The generation time was three years per generation, the mutation rate was 4.08 x 10^-9^ per nucleotide per generation.

## 3 RESULTS

### 3.1 Genome assembly and annotation

We initially assembled a draft genome from 16.7 Gbp of Nanopore and 21.4 Gbp of Illumina reads (Table S1). We produced 4,140 contigs and 607 Mb in total length with a contig N50 value of Mbp (Table S2). To enhance assembly contiguity, we generated Chicago and Hi-C libraries using Dovetail Genomics (Scott Valley, California), resulting in a genome size of 607 Mb across 2703 scaffolds with a contig N50 value of 24Mbp, with 23 chromosome-level assemblies. Blobtools analysis indicated that 94.97% (604 Mb) of the assembly aligns with the Chordata phylum, whereas 2.34% (2 Mb) did not match any known categories (Figure S1). The estimated genome size based on k-mer analysis is 604 Mb (Figure S2), which is consistent with the assembled size attributed to Chordata in the Blobtools analysis.

We obtained nine additional Gobioidei genomes from public sources (Table 1) to study Gobioidei evolution. To ensure the quality of further analyses, the selection criterion was that their genomic BUSCO scores must be higher than 90%. The BUSCO score for our assembled loach goby genome was 98.43%, the highest among the species compared (94.56% to 98.43%). Gobioidei species genome sizes, ranging from 607 to1343 Mb, appear to increase from basal to derived groups (Figure 1a). Notably, the loach goby was placed in the most basal position and has the smallest genome among the studied species.

BRAKER2.1.6 (Hoff et al., 2019) was used for genome annotation. We annotated 25,259 loach goby genes with 96.24% BUSCO completeness (Table 2 and Table S3). Among these genes, 89.84% match InterProScan, 66.62% have GO terms, 66.06% have orthologs in KEGG, and 48.85% are annotated across all three databases (Figure S3). On average, each mRNA comprises 10.5 exons that are approximately 160 bp long on average. These data are consistent with other annotated genomes (Table S3). However, the mean loach goby intron is only 892 bp long, considerably shorter than in other species (1020 to 2688 bp, p < 10^-3^; Table 2).

**Table 2.**
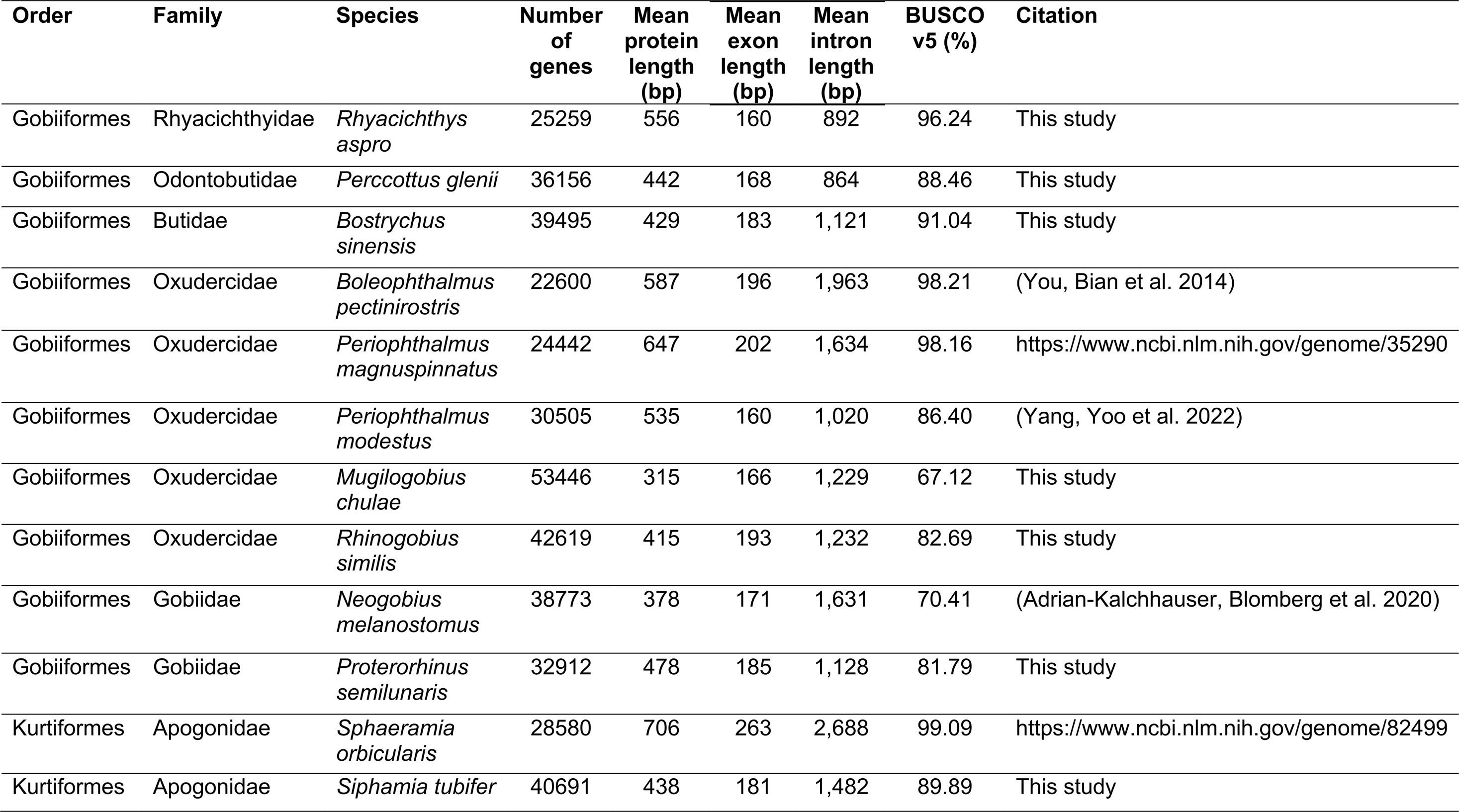
Statistics of the predicted gene.

To examine chromosomal evolution in Gobioid fishes, we used DAGchainer (Haas et al., 2004) to deduce synteny relationships between the loach goby genome and other chromosome-level assembled genomes, including *P. magnuspinnatus*, *P. modestus*, and *S. orbicularis*. We see one to one chromosome correspondence among these species, with the exception of chromosomes 12 and 23 of *P. magnuspinnatus* that matched the chromosome 12 in the other species. Thus, despite considerable genome size variability, Gobioid fish chromosome numbers remain remarkably conserved (Figure 2 and Table S4).

**Figure 2.**
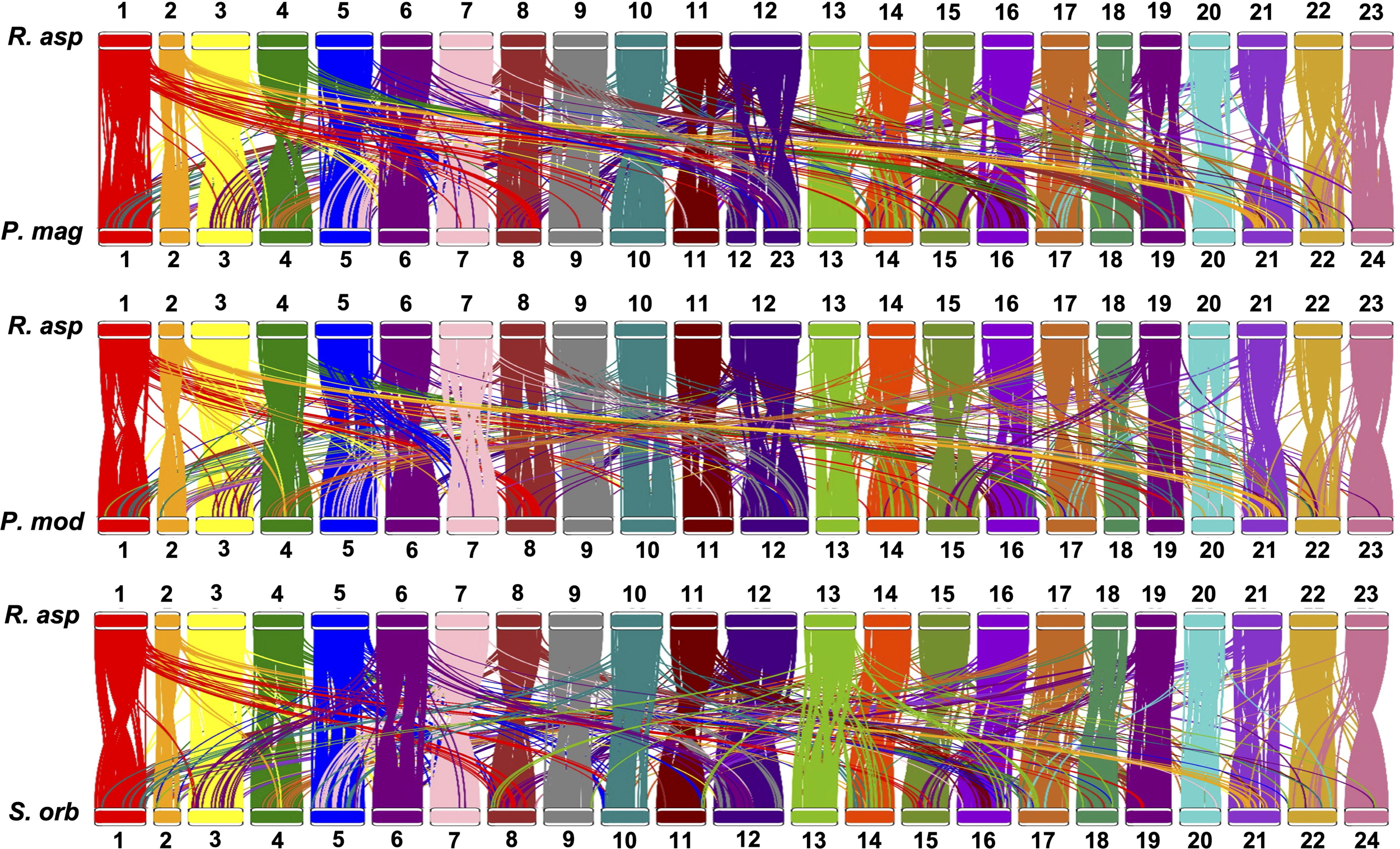
Syntenic chromosome conservation in gobioid fishes. The synteny relationships among *Rhyacichthys aspro* (R. asp), *Periophthalmus magnuspinnatus* (P. mag), *P. modestus* (P. mod), and *Sphaeramia orbicularis* (S. orb) revealed the conservation of chromosome numbers among gobioid fishes. Chromosomal orders referenced here are based upon the configuration in *P. modestus*. It is noteworthy that what is identified as chromosome 12 and 23 in *P. magnuspinnatus* correspond to what is designated as chromosomes 12 in other species. Additionally, *S. orbicularis* possesses only 23 chromosomes, lacking a distinct chromosome 23; instead, its chromosome 24 is homologous to chromosome 23 of *R. aspro* and *P. modestus*.

We identified 2100 single-copy orthologs in the ten sampled gobioid and two Kurtiformes genomes using Orthofinder (Emms & Kelly, 2019). These orthologous genes were concatenated and aligned to construct a maximum-likelihood phylogenetic tree (Figure 3). The phylogeny is consistent with previous studies, showing that Rhyacichthyidae and Odontobutidae together form the basal clade in Gobioidei (Agorreta et al., 2013; C. E. Thacker et al., 2023; Tornabene, Chen, & Pezold, 2013) followed by Butidae. Oxudercidae and Gobiidae are sister clades and are the most derived groups. Molecular dating was performed using MCMCtree (Z. Yang, 2007) and incorporated data obtained from TIMETREE (Kumar et al., 2022). The divergence between Gobiiformes and Kurtiformes was estimated at around 88 million years ago (MYA; 95% interval 86−92 MYA), at the lower end of previous estimates between 86 and 102 MYA (Alfaro et al., 2018; Hughes et al., 2018; Rabosky et al., 2018).

**Figure 3.**
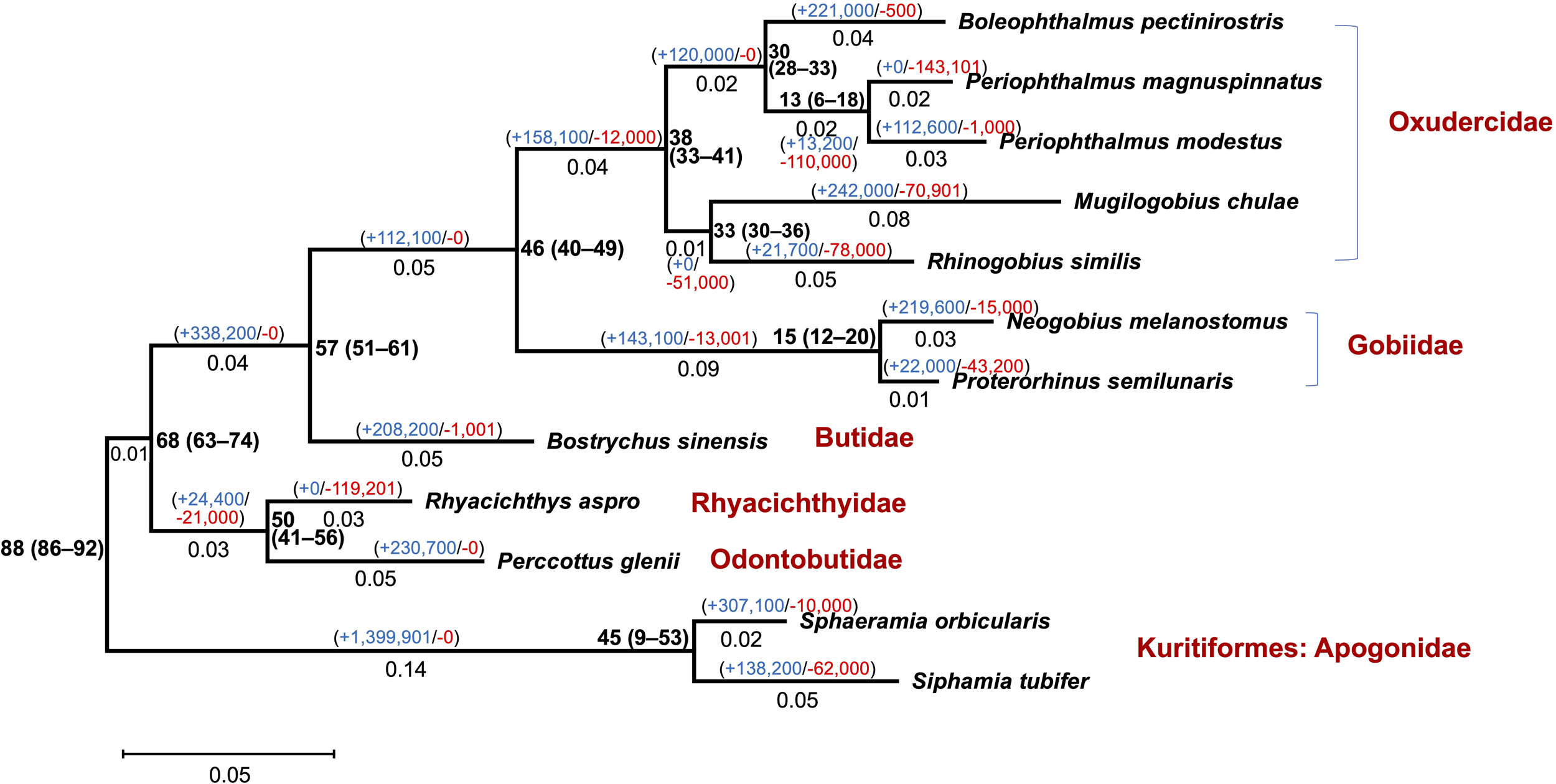
Phylogeny of Gobioidei. The phylogeny was constructed using 2100 single-copy orthologs, with the Kurtiformes serving as the outgroup. All nodes are supported by > 90% bootstrap values. Adjacent to the nodes, numbers represent the divergence times in millions of years, with the 95% confidence within parentheses. Below the branches, numbers indicate genetic distances, while gains (in blue) and losses (in red) of repeat elements are shown above the branches.

### 3.2 Repetitive elements and Gobioid evolution

Mirroring the trend in genome size, the proportion of repeat elements (REs) tends to increase from basal to derived Gobioidei groups. Notably, among all the species compared, the loach goby genome contains the fewest REs (Table 1). Furthermore, this holds for each individual RE type (Figure 4; Table S5a). Genome sizes are strongly positively correlated with RE (ρ =0.85, p< 0.01) after correcting for phylogenetic relationships using phylogenetically independent contrasts (PIC) (Felsenstein, 1985) (Figure S4a). Excluding REs strikingly reduces genome size variability, indicating that RE changes predominantly drive genome size variation (Figure S4b).

**Figure 4.**
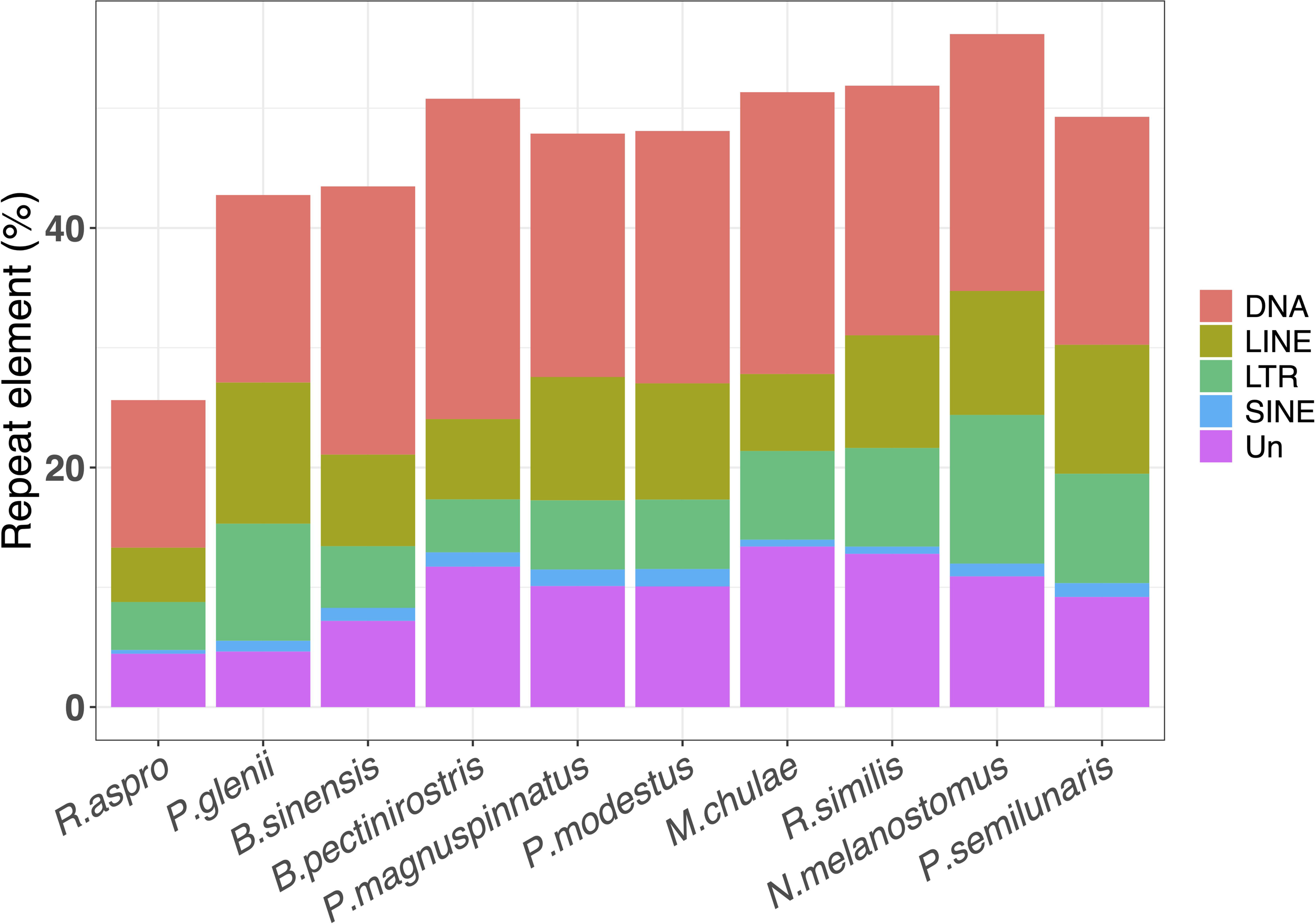
Proportions of different repeat elements in the Gobioidei genomes. DNA: DNA transposon; LINE: Long interspersed nuclear element; SINE: Short interspersed nuclear element; LTR: Long terminal repeat; UN: unknown.

The higher incidence of REs in the recently derived Gobioidei taxa may have resulted either from an acceleration of RE expansion or more active repetitive sequence reduction in the basal species. To distinguish between these possibilities, we used CAFÉ to analyze RE dynamics across the Gobioidei phylogeny (Figure 3). We see a general reduction of RE activity in the basal Gobioidei groups, indicating that genome enlargement in derived Gobioidei taxa is predominantly due to repetitive element expansion.

We constructed repeat landscapes to examine the evolutionary history of RE expansion (Figure 5) among our taxa. The loach goby genome has undergone less RE expansion than any of the other species examined. *Percccotthus glenii* (Odontobutidae) also shows limited historical RE expansion with a notable increase in recent activity (divergence level < 5%) (see also Figure 3). Butidae and Oxudercidae fishes have experienced a burst of repetitive element expansion leading to a high prevalence of among-locus divergence between 10− 20%. In contrast, Gobiidae species show a more recent onset of RE expansion (divergence level < 10%). In all cases, DNA transposons dominate RE expansion dynamics.

**Figure 5.**
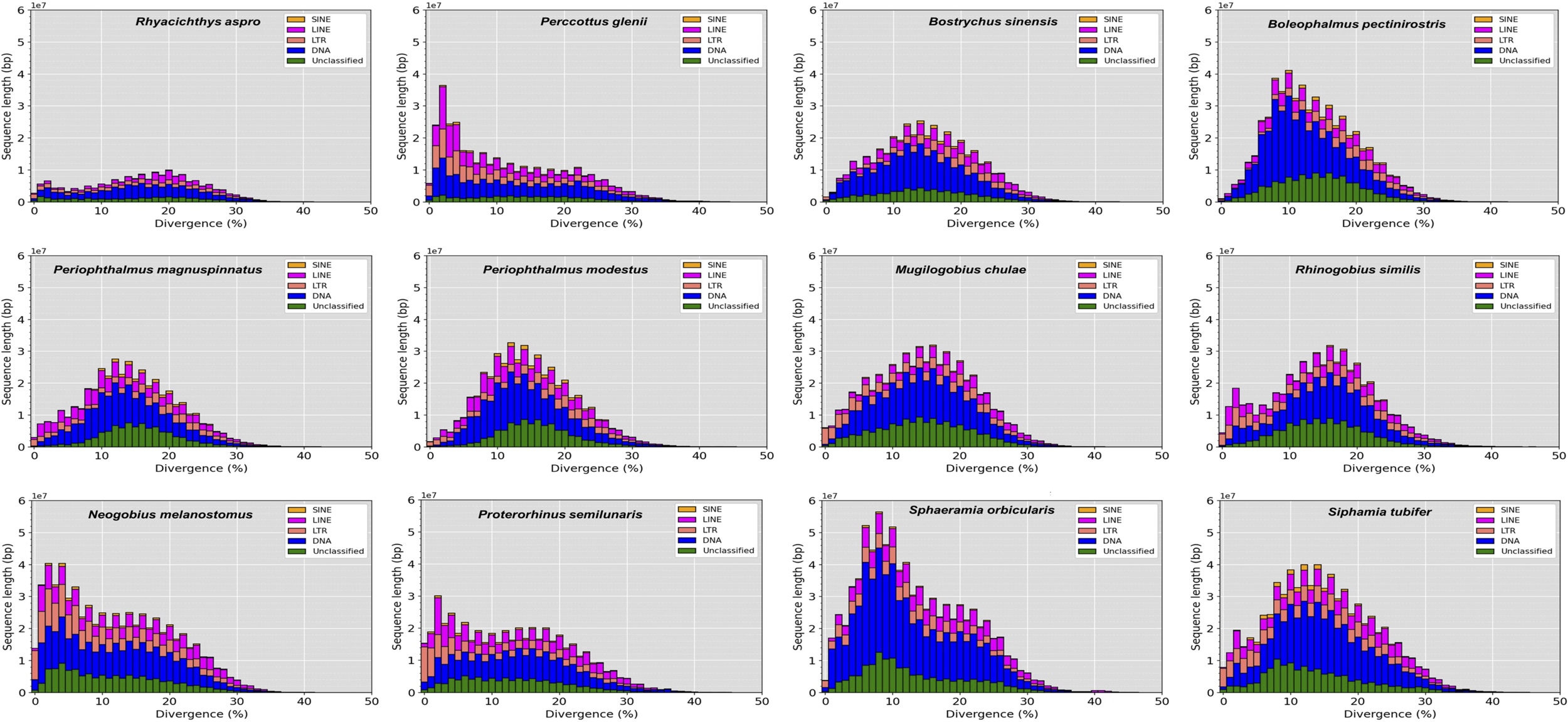
Comparison of the repeat landscapes among the 12 studied genomes. Note that the majority of expanded repeat elements are DNA transposons.

Figure 3 shows that branches leading to clades harboring few repetitive elements, including Rhyacichthyidae, Odontobutidae, and Butidae, are shorter than those leading to RE rich groups (Oxudercidae and Gobiidae). Indeed, the loach goby branch is the shortest. To investigate this pattern, we computed genetic distances between Gobioidei fishes and the Kurtiformes outgroup (Table S6). The genetic distances between the Gobioidei species and the Kurtiformes, which diverged from a shared ancestor concurrently, can be utilized as proxies for their mutation rates, as the discrepancies in these distances reflect the differential mutation rates that have occurred post-divergence. We see strong positive correlation of genetic distance and RE prevalence (ρ = 0.87, p < 0.01, corrected for phylogeny using PIC; Figure 6). This correlation is found at both synonymous (ρ = 0.75, p < 0.02; Figure S5a) and nonsynonymous sites (ρ = 0.9, p < 0.01; Figure S5b). When different types of RE are evaluated separately, genetic distances are positively correlated with DNA transposon (ρ = 0.85, p < 0.01) and unclassified RE abundance (ρ = 0.87, p< 0.01; Figure 6), but not with retrotransposon incidence (i.e., SINEs, LINEs, LTRs, and all retroelements together; Figure S6). Among different DNA transposon superfamilies, the correlations persist with *hAT* (ρ = 0.68, p = 0.05) and *hAT.Ac* transposons (ρ = 0.73, p = 0.03; Table S5b, Figure S7). DNA repair after transposon excision and insertion can result in an increased mutation rate of adjacent sequences (Wicker et al., 2016). Therefore, in addition to looking at transposon abundance, we examined the cumulative number of alterations associated with RE activity (insertions and excisions). We still see strong correlations of these more comprehensive metrics with genetic distances for all DNA transposons (ρ = 0.92, p< 0.01) and *hAT* transposons in particular (ρ = 0.87, p< 0.01; Figure S7). Notably, genetic distances of mitochondrial genes exhibit no association with RE content (Table S7, Figure S8).

**Figure 6.**
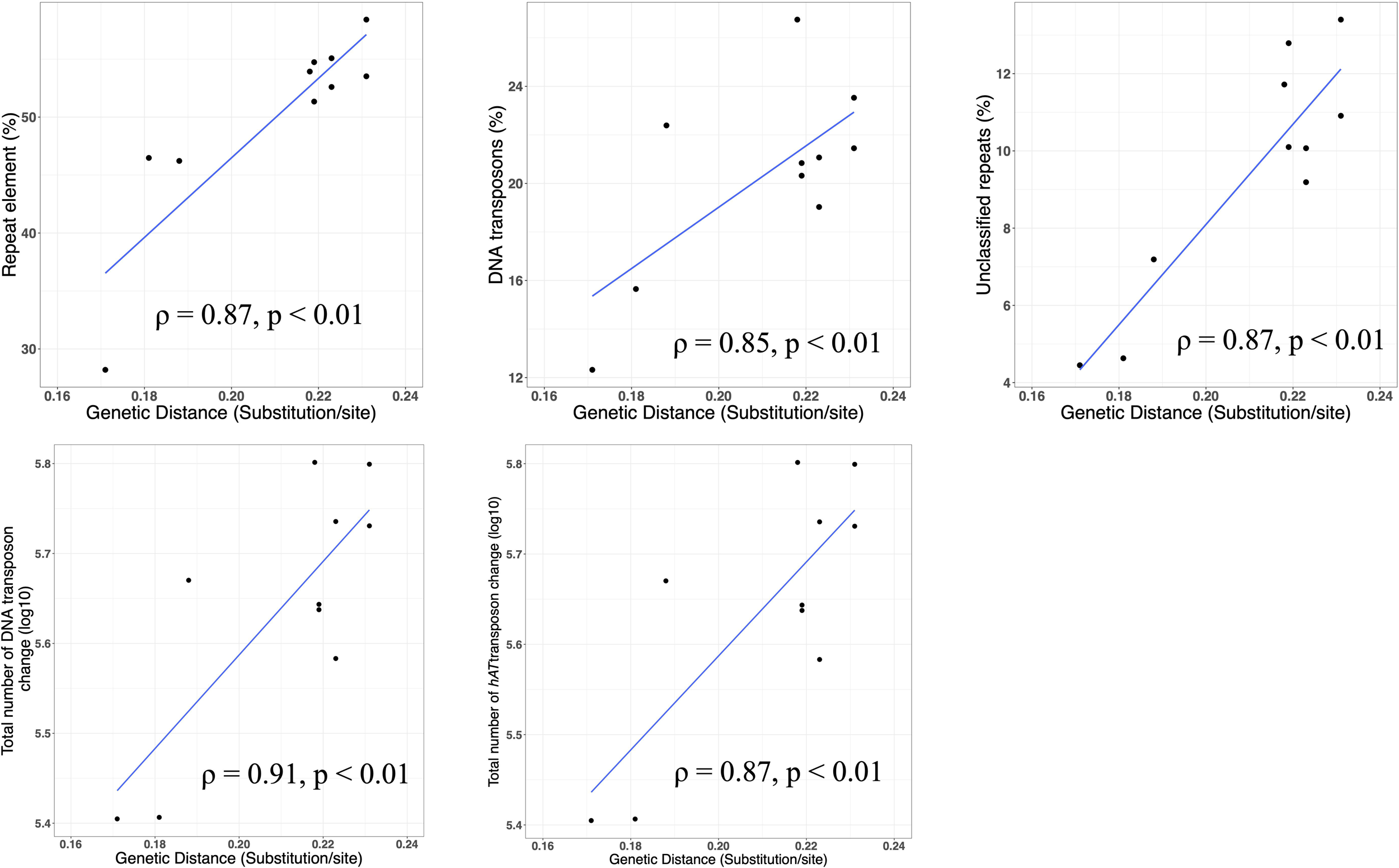
Correlation between different repeat element types and mutation rate. The genetic distances were distance to outgroup Kurtiformes. Since all Gobioidei were derived from the outgroups at the same time, the genetic distances are served as a proxy for mutation rate. The correlation was conducted using Kendall’s ρ coefficient after correcting for phylogeny using phylogenetically independent contrasts (see text for details)

We next examine RE prevalence in coding sequences and regulatory regions (within 5Kb upstream of translation start site), as repetitive elements within these regions can influence gene function and lead to phenotypic diversity (Chuong, Elde, & Feschotte, 2017; Fotsing et al., 2019; Shi et al., 2023). DNA transposons are significantly more abundant in above two regions than retrotransposons (SINEs, LINEs, and LTRs) (p < 0.01; Wilcoxon rank-sum test; Figure 7). While the differences in percentages may appear marginal, the number of DNA transposons is much larger than that of various types of retrotransposons. Therefore, even small percentage differences reflect considerable variations in actual counts, which could have important consequences for genomic diversity and evolution. This enrichment is mostly due to excess prevalence of *hAT* and *hAT.Ac* DNA transposons (p < 0.01). The difference between DNA transposons and retrotransposons is less pronounced in Kurtiformes.

**Figure 7.**
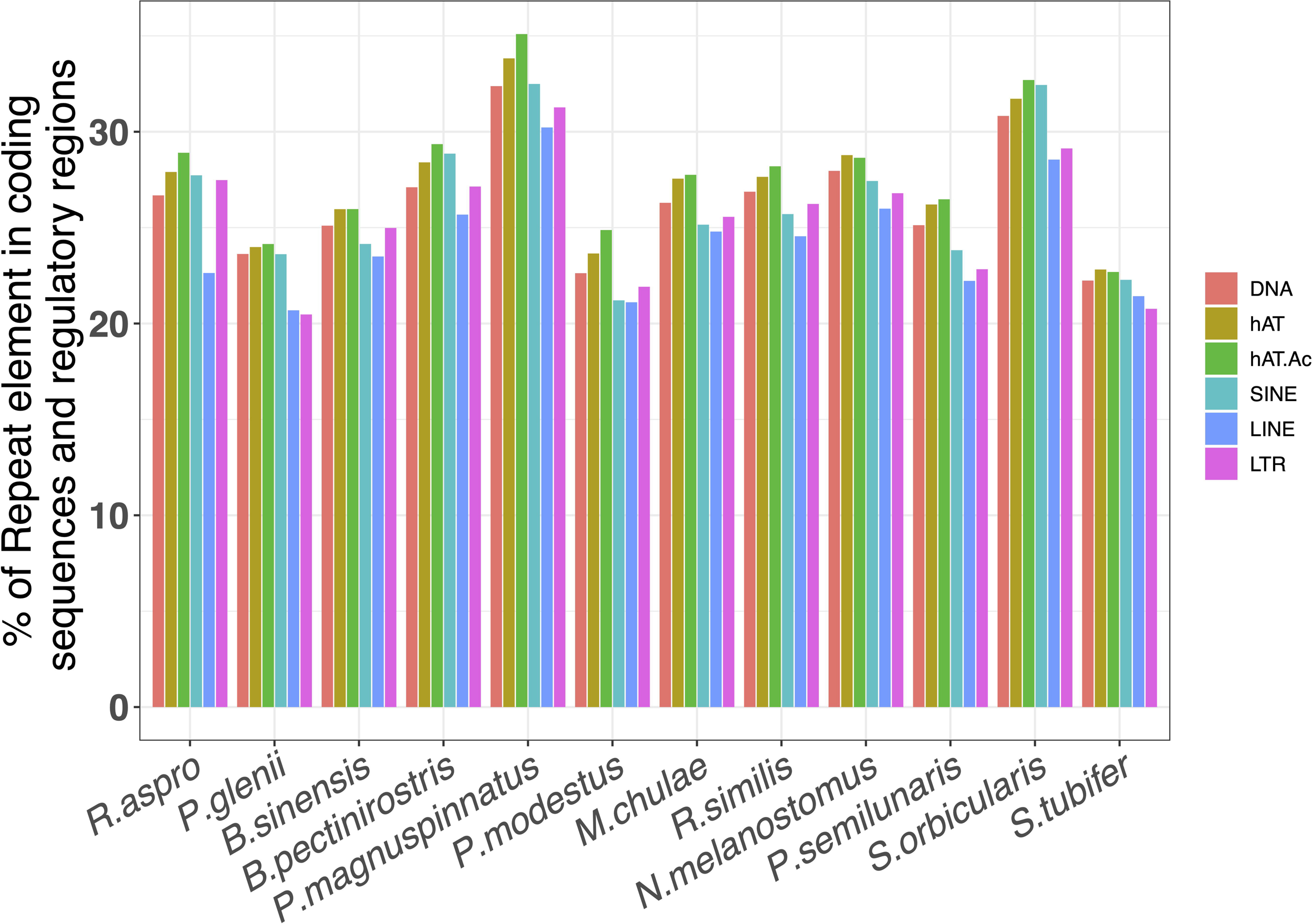
The proportion of various types of repeat elements in functional regions. The functional regions include coding sequences and the regions within 5 kilobases (Kb) directly upstream of the translation start site

### 3.3 Genes related the loach goby natural history

Our analyses of the loach goby genome also shed light on the special features of this species (Table 3, Table S8-S13). Table S8-S13 listed the expanded/ reduced GO term (Table S8-S10), protein family (Table S11), protein domain (Table S12) and KEGG number (Table S13). Specifically, the loach goby has fewer genes associated with neuromast development and differentiation than other Gobioidei species, including genes involved in posterior lateral line neuromast primordium migration (GO:0048920) (Table S14a) and anterior lateral line neuromast development and differentiation (GO:0035676; GO:0048901, GO:0048903) (Table S14b). These differences may contribute to the unique mechanosensory lateral line system observed in the loach goby (see Introduction). Some genes in these GO terms are known to be involved in neuron system development and function. For example, sphingosine-1-phosphate receptor 2 (s1pr2) (Table S14a) has an important role in the development of otic vesicles, neuromasts, and survival of hair cells. Misregulation of s1pr2 leads to change in hair cell and neuromast abundance and their lateral line positioning in zebrafish (Z. y. Hu et al., 2013). The insulinoma-associated protein 1a (insm1a) (Table S14b) is a zinc-finger transcription factor playing a variety of roles in cell formation and differentiation of vertebrate central and peripheral nervous systems (Gong et al., 2017). Knockdown of insm1a expression in zebrafish leads defects in primordium migration, resulting in fewer neuromasts (He et al., 2017).

**Table 3.**
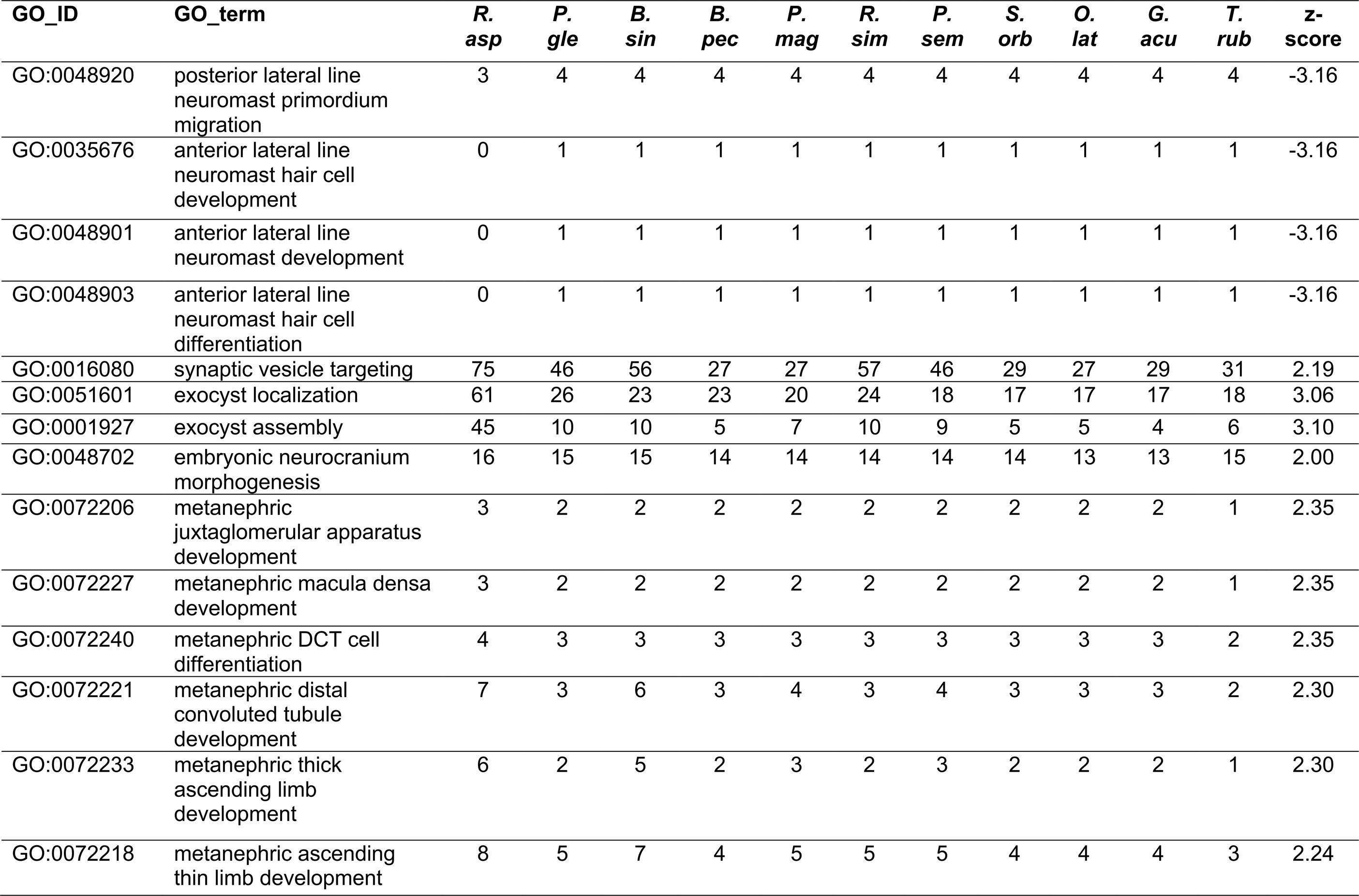
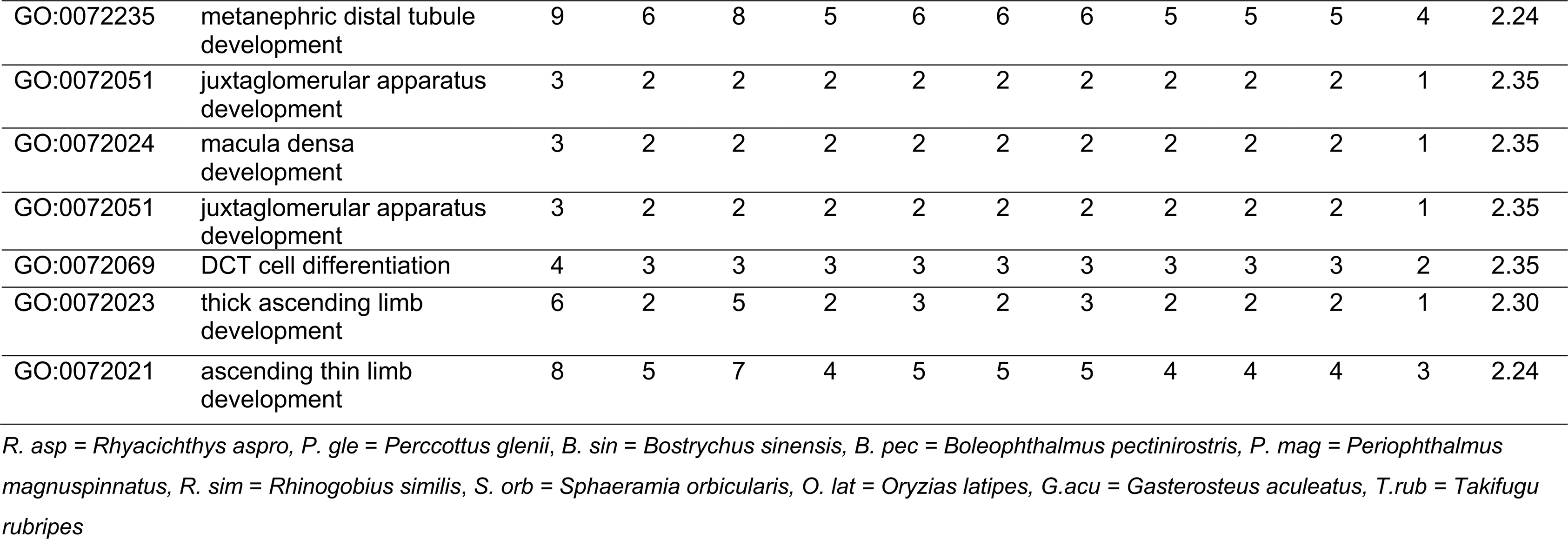
Significantly enriched/ depleted GO terms biological process in *Rhyacichthys aspro*.

Interestingly, the loach goby genome also has an overabundance of genes involved in synaptic formation and in sensory information processing, including synaptic vesicle targeting (GO:0016080), and exocyst localization (GO:0051601) and assembly (GO:0001927). Consequently, while loach goby has a primitive lateral line system, it might have evolved unique sensory perception and lateral line development pathways at the molecular level.

Morphologically, loach goby is characterized by a flattened head and abdomen These characteristics may be critical for adaptation to torrential currents. It is thus notable that the genome of this species is enriched in genes involved in embryonic neurocranium morphogenesis (GO:0048702) (Table 3, Table S14c), suggesting a genetic foundation for these morphological adaptations. For example, we find an additional copy of the protein transport factor Sec23A and transforming growth factor beta-2 proprotein (Tgfb2). Mutations in Sec23A are associated with facial dysmorphisms and late-closing fontanels in humans (Boyadjiev et al., 2011). Tgfb2 is a precursor of transforming growth factor β-2 (TGF-β-2) that regulates fetal rat cranial suture morphogenesis (Opperman, Adab, & Gakunga, 2000).

Loci involved in metanephric distal tubule (GO:0072235) and metanephric ascending thin limb development (GO:0072218) may underly the loach goby life history traits. Loach goby larvae drift out into the open ocean and return to fresh water only after becoming juveniles. Therefore, osmoregulation is expected to be particularly important. Some genes, such as POU Domain Class 3 Transcription Factor 3 (POU3F3) and deleted in malignant brain tumors 1 (DMBT1) (Table S14d), in these GO categories are involved in kidney development and osmoregulation. For example, an amino acid mutation in mouse POU3F3 causes reduced nephron number and impaired kidney development (Rieger et al., 2016). Deletion of DMBT1 results in distal renal tubular acidosis, a kidney condition that impairs removal of acids from blood into urine (Gao et al., 2010).

To identify genes that might underly loach goby adaptive evolution, we used Tajima’s D test to identify selective sweeps. We find 1296 genes with negative Tajima’s D values (Table S15). Several GO categories are overrepresented among them (Table S16). In particular, synapse organization (GO:0050808) and calmodulin binding (GO:0005516) may influence sensory perception. The organization of synapses plays a pivotal role in transmitting and processing sensory information across neural circuits (Balaskas, Abbott, Jessell, & Ng, 2019). Calmodulin plays a significant role in sensory systems, including vision, hearing, taste, and touch, by mediating the signaling pathways involved in detecting and processing sensory stimuli (Fain, Hardie, & Laughlin, 2010; Gillespie & Muller, 2009; Wu, Lewis, & Grandl, 2017; Zhang et al., 2003). Using Phenotype Enrichment Analysis (Table S17) (Weng & Liao, 2017), we also found that genes putatively under selective sweeps (Table S15) are disproportionally involved in cranial cartilage development. This finding, coupled with the observed enrichment of genes linked to embryonic neurocranium morphogenesis (GO:0048702) (Table 3) in the loach goby genome, implies a genetic basis for the species’ distinctive flattened head and abdomen. These morphological traits are likely adaptive and facilitate survival in the fast-flowing waters of their native habitat.

### 3.4 Population demography

To understand loach goby genetic diversity and population demography, we also generated four additional loach goby genomes (Table S18). Their heterozygosity ranges from 0.34% to 0.37% with an average nucleotide diversity π of 4.12×10^-3^ (or one heterozygous single nucleotide polymorphism every 233 bp).

Our analysis using Multiple Sequentially Markovian Coalescent 2 (MSMC2) (Schiffels & Wang, 2020) suggests that loach goby populations have experienced an expansion around 5 x 10^5^ years ago (Figure 8a). The population size grew to 4 x 10^5^ approximately 10^5^ years ago, and then to 1 x 10^6^ by 10^4^ years ago. However, it is worth noting that the MSMC2 tends to be unstable with large haplotype sizes (hap = 10) and may not provide accurate estimates of recent history (X. Liu & Fu, 2020). Therefore, we also utilized Stairway Plot 2 for a more recent demographic estimation (X. Liu & Fu, 2020). Our estimates suggest that loach goby populations began to grow around 6 x 10^5^ years ago, with the size hovering around 4 x 10^5^ between 400,000 and 10,000 years ago (Figure 8b). The populations then experienced a decline, diminishing to an estimated 100,000 individuals, which corresponds to its current population size.

**Figure 8.**
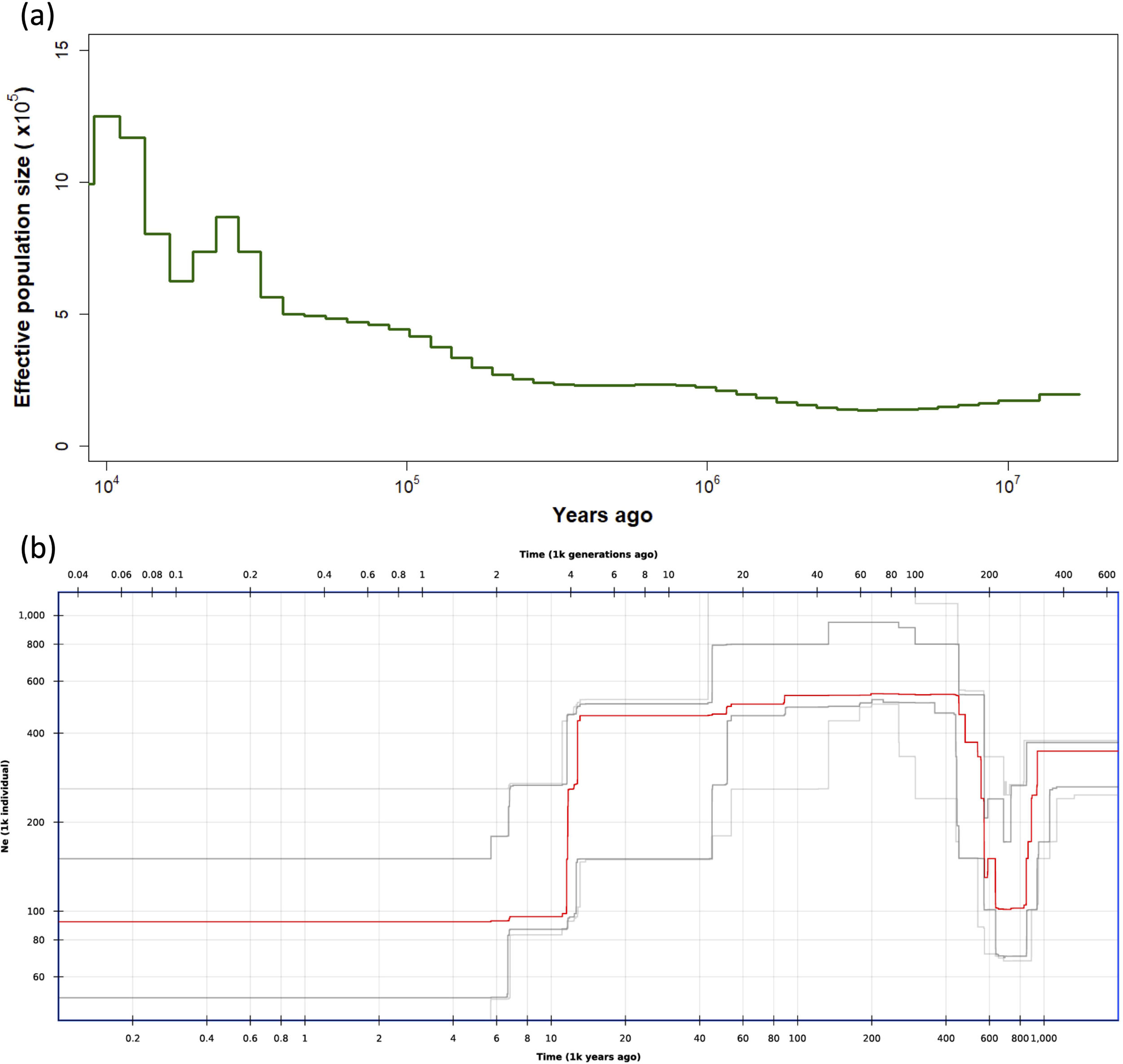
Population demography of loach goby by MSMC2 (a) and stairway plot 2 (b). The demography was based on the genome sequences of five loach gobies.

This recent demographic decline is strongly associated with sea-level change. The peak in loach goby numbers coincided with lower sea levels and declined as levels rose at the close of the last glacial period, about 10,000 years ago. These patterns may be linked to change in habitat availability, since higher sea levels probably shrink freshwater habitats required by the species in their juvenile and adult stages.

## 4 DISCUSSION

### 4.1 The loach goby genome elucidates Gobioidei diversification

The loach goby genome is small and has a low mutation rate, likely due to a relatively light repetitive element (RE) load. While chromosome numbers are remarkably conserved (Figure 2) across Gobioidei taxa, RE content varies widely, ranging from 28 to 60% of the genome. There is a marked increase in both genome size and repeat element, particularly DNA transposon, prevalence as we go from basal to derived taxa. DNA transposons account for 12% to over 26% of the genomes and 30 to 50% of REs. DNA TEs also predominate among expanding classes of repeats (Figure 5). Furthermore, these expansions occur mostly in the species rich derived families, suggesting a positive correlation between RE proliferation and species diversification.

We also see that DNA transposons, especially the *hAT* and *hAT.Ac* families, are disproportionally found in coding and regulatory regions (Figure 7), consistent with previous studies (Munoz-Lopez & Garcia-Perez, 2010; Wicker et al., 2016; Woodard et al., 2012) We further see a strong positive correlation between transposon activity (insertion and excision) and level of divergence, suggesting that RE movement causes the elevated mutation rates in adjacent sequences (Wicker et al., 2016). Since these transposons are prevalent in functional regions, their mutations likely lead to new functional variants, thereby enhancing the functional complexity required for adaptation to diverse environmental conditions (Chuong et al., 2017; Fotsing et al., 2019; Shi et al., 2023). Therefore, the rapid proliferation of DNA transposons may be a contributing factor to the rapid diversification observed within Gobioidei.

RE activity can also facilitate speciation by causing large-scale genomic variation that creates interspecific genetic incompatibilities (Serrato-Capuchina & Matute, 2018). High RE prevalence in mammalian genomes is associated with increased speciation rate (Ricci, Peona, Guichard, Taccioli, & Boattini, 2018). It is thus possible that the diversification of Gobioidei, one of the most species rich vertebrate orders, was likely facilitated by the accelerated modification and expansion of REs in their genomes (Figure 9), promoting novel gene combinations, duplications, and losses, as well as altered gene functions and regulatory networks. These genome-wide changes in turn may have enhanced the capacity of these taxa to adapt to diverse aquatic ecosystems.

**Figure 9.**
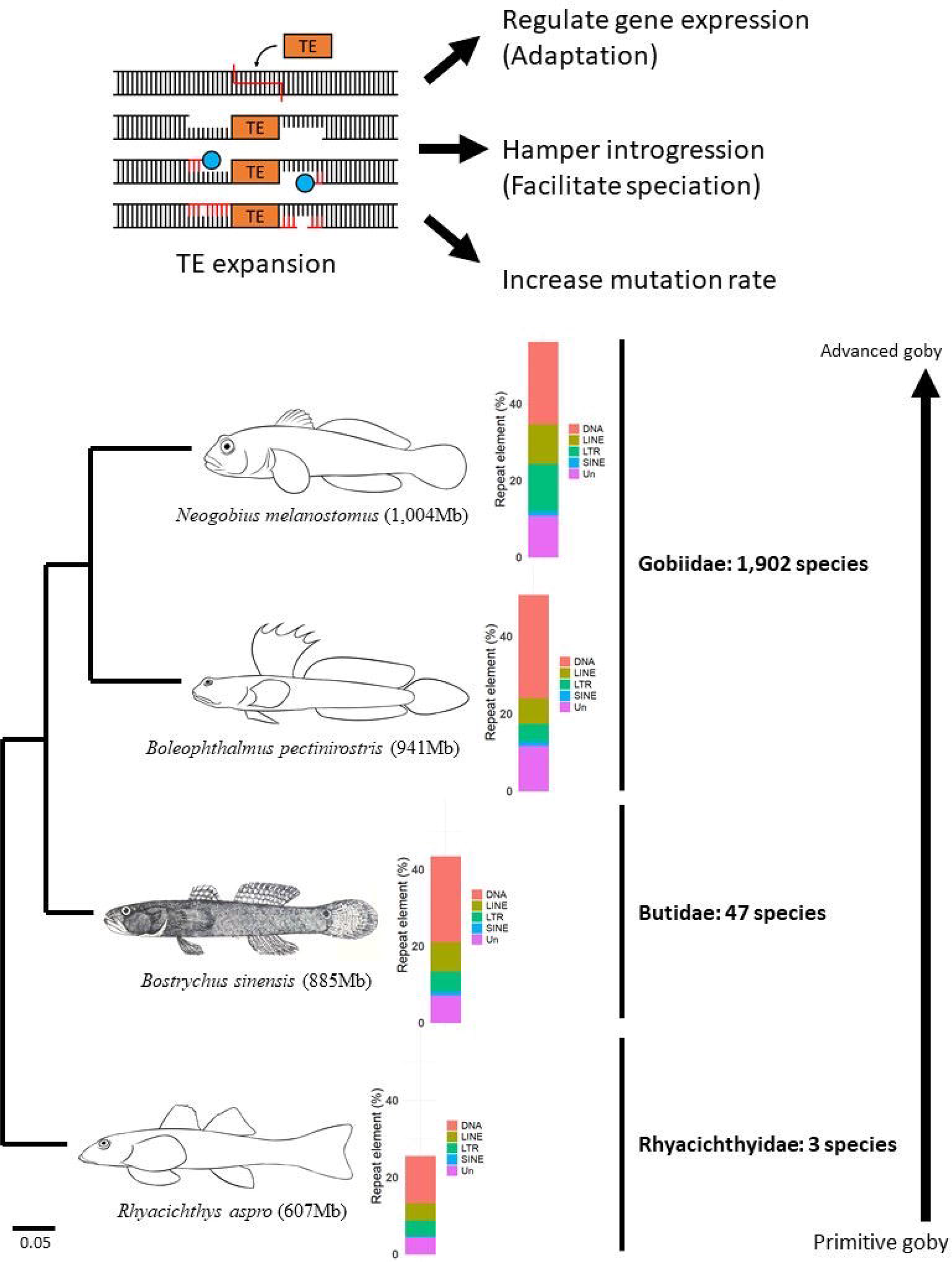
Schematic illustration of Gobioidei evolution. Within this suborder, there is a marked increase in both genome size and repeat element, particularly DNA transposon, from basal to more derived species. The expansion of DNA transposons is linked to increased mutation rates which may amplify functional complexity for adaptation to varied environments and drive speciation through extensive genomic variations that establish genetic incompatibilities between species. This expansion, notably within functional regions, likely contributes to the remarkable diversification and ecological adaptation observed in this diverse group of vertebrates.

### 4.2 Genomic signatures of adaptation

Analyses of the loach goby genome also reveal special adaptive features of this species. The gnome is relatively poor in genes associated with lateral line and neuromast development, but is rich in loci related to synaptic development (Table 3). This suggests that despite the presence of the lateral line system that is primitive among Gobioidei, this species may have involved a unique molecular approach to sensory perception and lateral line development. Genes putatively under positive selection are also disproportionally involved in synapse organization (GO:0050808) and calmodulin binding (GO:0005516) (Table S16), also likely related to the sensory perception (Balaskas et al., 2019; Fain et al., 2010; Gillespie & Muller, 2009; Wu et al., 2017; Zhang et al., 2003).

The loach goby genome has copies of Sec23A and Tgfb2 (Table S14c) not present in related gobioid species. These loci are involved in embryonic neurocranium morphogenesis (GO:0048702) (Table 3). Mutations on these genes affect cranial development and suture morphogenesis (Boyadjiev et al., 2011; Opperman et al., 2000). Cranial cartilage development is also a category overrepresented among putatively adaptively evolving genes (Table S17). These loci may be associated with flattened heads and abdomens that characterize loach gobies, phenotypes that may underly adaptation to strong currents. Interestingly, we also found that the genome of a hill-stream loach *Beaufortia kweichowensis* from the Balitoridae (Cypriniformes) family that also has a flattened cranium (Deng et al., 2021) adaptive in fast mountain streams is also enriched in the GO: GO:0048702 category (Table S19a). The *B. kweichowensis* genome contains a paralog of Sec23A, Sec23B, that is also evolved in embryonic neurocranium morphogenesis (Lang, Lapierre, Frotscher, Goldenring, & Knapik, 2006).

Genes related to renal development and osmoregulation (Table 3 and Table S14d) are also prevalent in the loach goby genome, suggesting a molecular mechanism of adaptation to its amphidromous life history. To demonstrate that changes in osmoregulation are particularly important in fish that move between marine and freshwater environments, we also analyzed the genome of the European eel (*Anguilla anguilla*) that is a typical catadromous fish (van Ginneken & Maes, 2005). Its genome also shows a strong enrichment in genes involved in osmoregulation (Table S19b).

## 5 CONCLUSION

The loach goby genome provides compelling evidence that repeat element dynamics drive Gobioidei adaptation and diversification against the background of chromosome number conservation (Figure 9). Specifically, the expansion of DNA transposons is linked to increased mutation rates, likely contributing to the rich diversity and adaptive capacity of the suborder. Our results suggest that RE-driven genetic alteration may be a driving force behind speciation and ecological success of one of the most species rich vertebrate orders, particularly in their adaptation to various aquatic ecosystems.

The loach goby genome also points to molecular mechanisms underlying unique sensory and morphological traits. The presence of an expanded repertoire of genes associated with synaptic development and sensory perception, alongside specific gene duplications related to cranial structure, underscores the loach goby’s evolutionary response to its ecological niche. These findings not only improve our understanding of the adaptive strategies employed by this species, but also underscore the broader evolutionary mechanisms that enable species to thrive in diverse and changing habitats. Comparative genomic analyses also reveal parallel evolution across species, reinforcing the importance of genetic adaptation through osmoregulation for aquatic species with complex life histories.

## AUTHOR CONTRIBUTIONS

The following are the authors’ contributions: designed research: TYW, and HYW; performed sampling: TYW, TYL, SPH and FYW; performed molecular experiment: TYW and FYW; performed bioinformatic analyses: HJL, YWW and HYW; visualization: HJL, CTT and HYW; statistical analyses: HJL and HYW; supervision: TYW, CTT, SMC and HYW; Writing–original draft: TYW, HJL, YWW, SMC and HYW; and Writing–reviewing–editing: TYW, HJL, YWW, SPH, FYW, CTT, SMC and HYW.

## ACKNOWLEDGEMENT

This study was funded by the National Science and Technology Council (MOST 102-2311-B-001-019 and MOST 108-2621-B-001-002 to TYW; MOST 109-2311-B-002-023-MY3 to HYW) and part of Taiwan BioGenome Project, funded by Academia Sinica, Taiwan (AS-Grant 23-23 to SMC).

## DATA AVAILABILITY AND BENEFIT SHARING STATEMENT

The loach goby (*R. aspro*) genome project was deposited at NCBI under BioProject No. PRJNA1010998. The Illumina DNA sequencing data were deposited under NCBI Accession No. SRR28458574; the Nanopore DNA sequencing data were deposited under NCBI Accession Nos. SRR28439223; the Hi-C sequencing data were deposited under NCBI Accession Nos. SRRXXXXXX. The Illumina RNA sequencing reads for different tissues were available under accession number SRR28461528 (eye), SRR28466848 (nose), SRRXXXXXX (brainSUB14339590), SRR28462307 (gill), SRR28466864 (muscle) and SRR28466849 (liver). The genome assembly with gene and transcript annotations has been deposited into GenBank under the accession number GCA_036850595.1 (ASM3685059v1).

## Ethics approval consent to participate

All animal experiments in this study were performed in accordance with guidelines of the animal ethics committee and were approved by the Institutional Animal Care and Use Committee (IACUC 20-12-1593), Academia Sinica.

