## Supplement Figures for "Chromosome-level genome assembly of the primitive loach goby, *Rhyacichthys aspro*, reveals mechanisms underlying Gobioidei diversification"

**Table of Contents (see supplemental excel file):**

**Table S1 - Description of sequencing data produced in this study**

**Table S2. Statistics of chromosome assembly.**

**Table S3. Statistics of the predicted gene.**

**Table S4a. Chromosome list in synteny.**

**Table S4b. Sequence accession number in GenBank.**

**Table S5a. List of repetitive sequence types**

**Table S5b. List of DNA transposons**

**Table S6a. The nucleotide diversity of single-copy genes of selected species.**

**Table S6b. The pairwise dS of single-copy genes of selected species.**

**Table S7. The pairwise genetic distance of mitochondria coding genes of selected species.**

**Table S8a. List of loach goby expanded GO term for biological process**

**Table S8b. List of loach goby reduced GO term for biological process**

**Table S9a. List of loach goby expanded GO term for molecular function**

**Table S9b. List of loach goby reduced GO term for molecular function**

**Table S10a. List of loach goby expanded GO term for cellular component**

**Table S10b. List of loach goby recuced GO term for cellular component**

**Table S11a. List of loach goby expanded protein family**

**Table S11b. List of loach goby reduced protein family**

**Table S12a. List of loach goby expanded protein domain**

**Table S12b. List of loach goby reduced protein domain**

**Table S13a. List of loach goby expanded KEGG number.**

**Table S13b. List of loach goby reduced KEGG number.**

**Table S14a. Genes include in GO:0048920.**

**Table S14b. Gene includes in GO:0035676, GO:0048901 and GO:0048903.**

**Table S14c. Genes include in GO:0048702.**

**Table S14d. Genes include in GO:0072235.**

**Table S15. List of genes with negative Tajima's D**

**Table S16. GO enrichment of genes with negative Tajima's D**

**Table S17. List of genes related to cranial cartilage development**

**Table S18. Mapping rate and coverage of the loach goby.**

**Table S19a. Gene expansion of *Beaufortia kweichowensis* in embryonic neurocranium morphogenesis.**

**Table S19b. Gene expansion of *Anguilla anguilla* in kidney-related GO.**

**Supplemental Figures.**

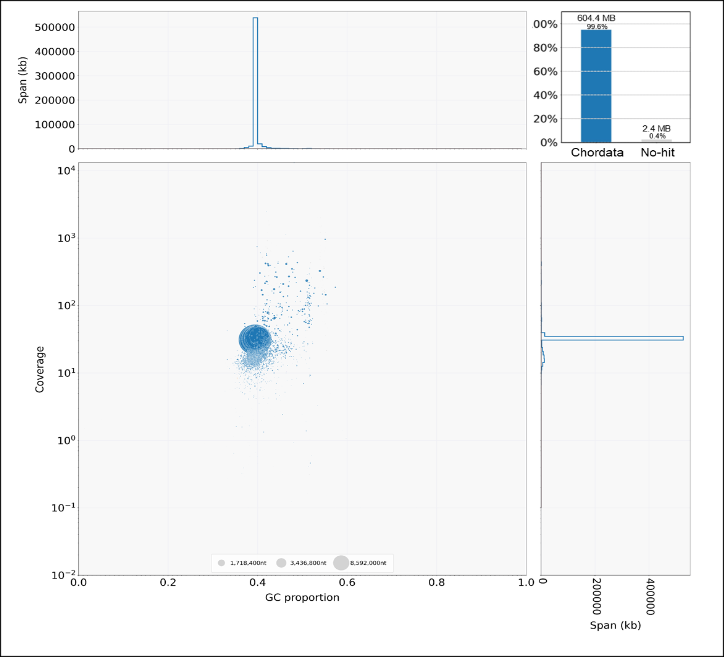

Figure S1. Blobplot result of the loach goby draft genome assembly.

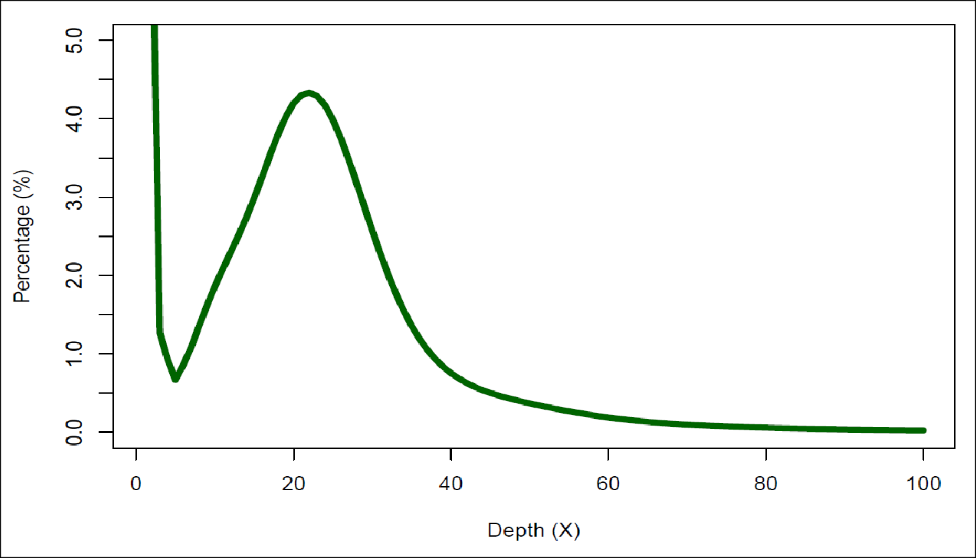

Figure S2. K-mer (k = 17) analysis for estimating the genome size of loach goby, *Rhyacichthys aspro*.

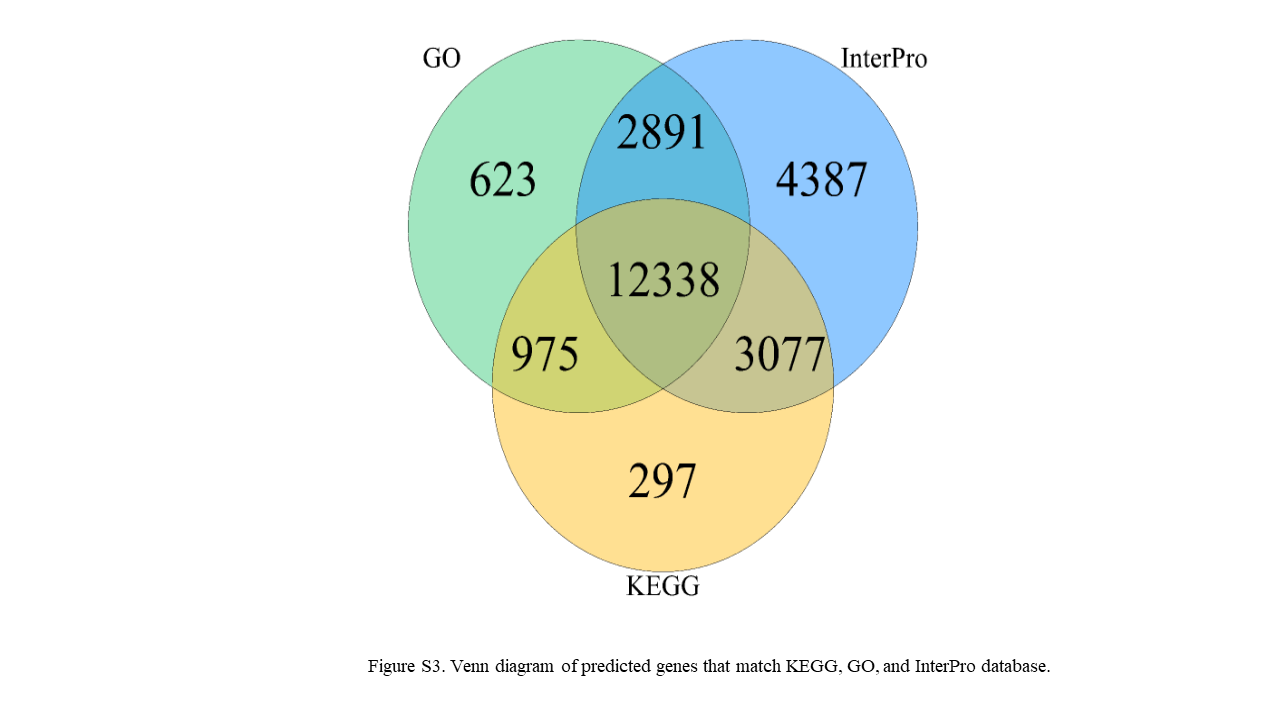

Figure S3. Venn diagram of predicted genes that match KEGG, GO, and InterPro database.

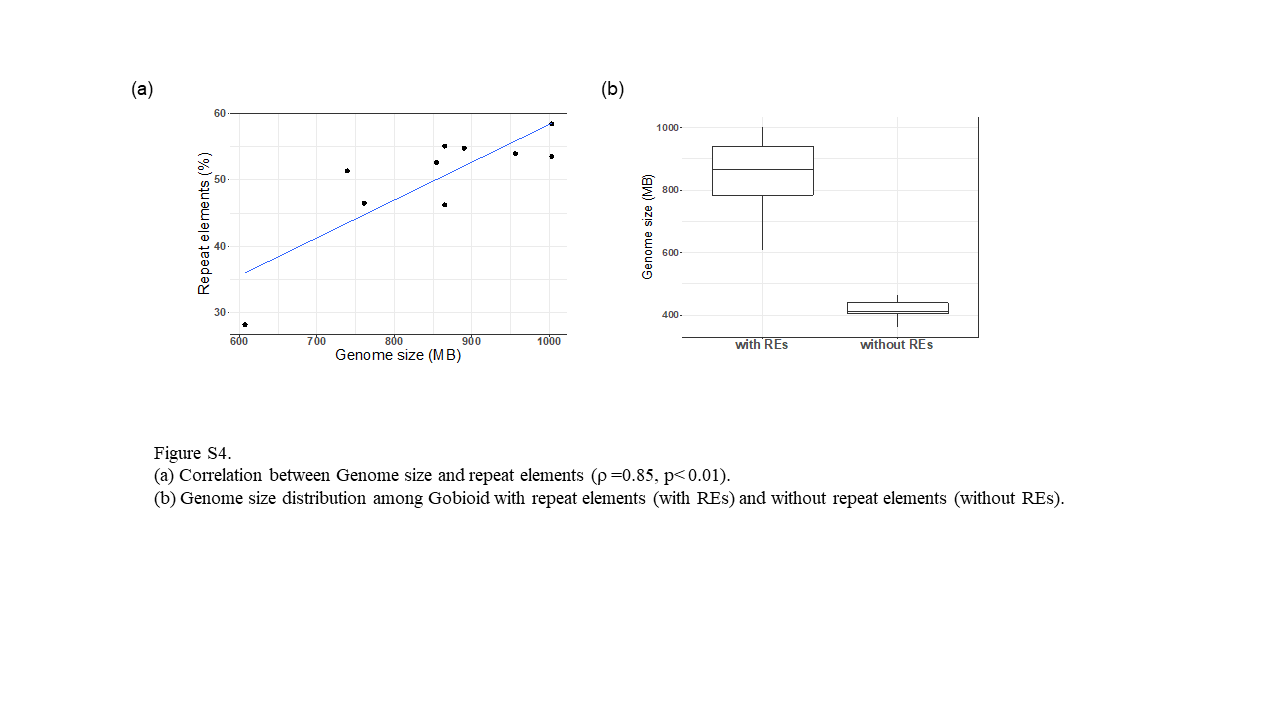

Figure S4. (a) Correlation between Genome size and repeat elements (ρ =0.85, p< 0.01); (b) Genome size distribution among Gobioid with repeat elements (with REs) and without repeat elements (without REs).

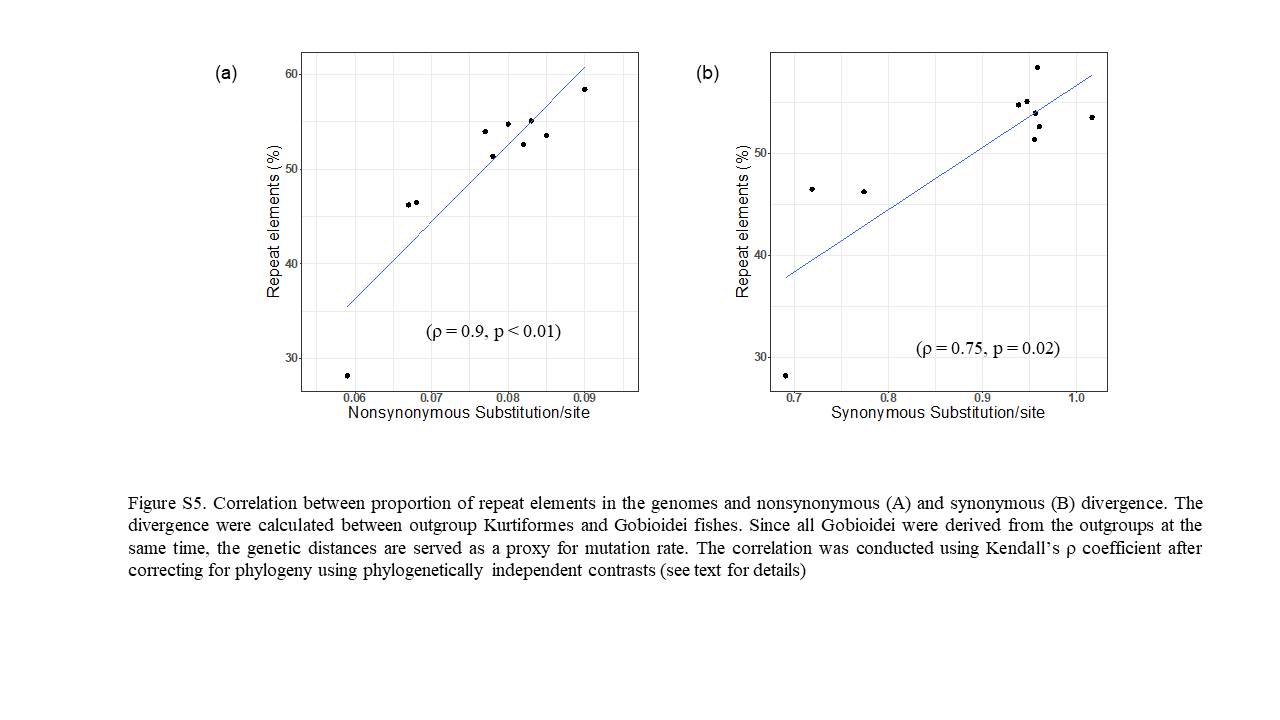

Figure S5. Correlation between proportion of repeat elements in the genomes and nonsynonymous (a and synonymous (b) divergence. The divergence were calculated between outgroup Kurtiformes and Gobioidei fishes. Since all Gobioidei were derived from the outgroups at the same time, the genetic distances are served as a proxy for mutation rate. The correlation was conducted using Kendall’s ρ coefficient after correcting for phylogeny using phylogenetically independent contrasts (see text for details)

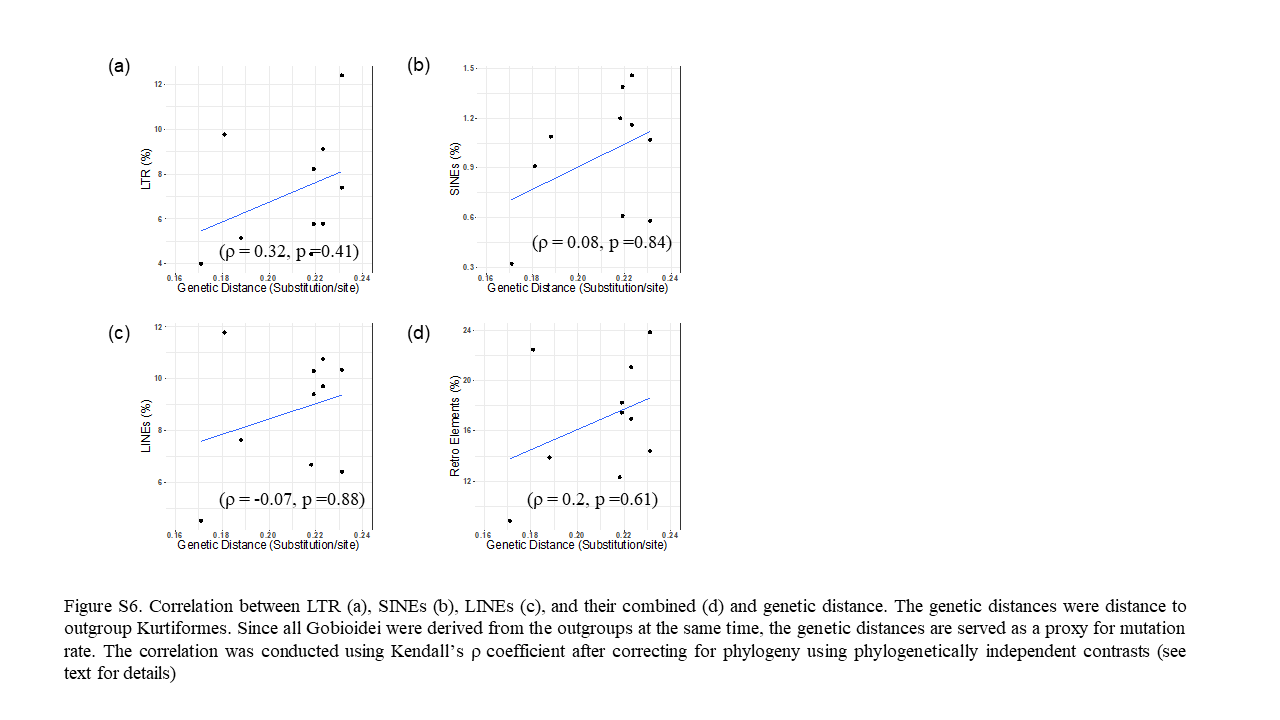

Figure S6. Correlation between LTR (a), SINEs (b), LINEs (c), and their combined (d) and genetic distance. The genetic distances were distance to outgroup Kurtiformes. Since all Gobioidei were derived from the outgroups at the same time, the genetic distances are served as a proxy for mutation rate. The correlation was conducted using Kendall’s ρ coefficient after correcting for phylogeny using phylogenetically independent contrasts (see text for details)

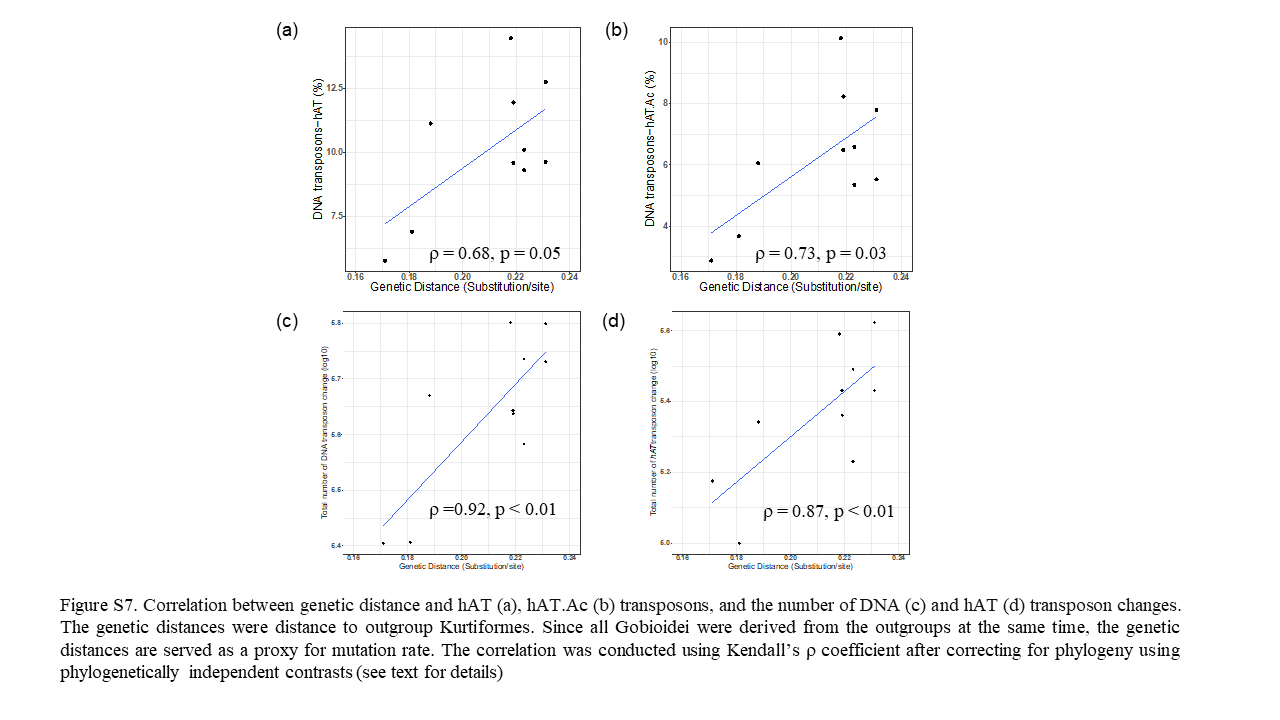

Figure S7. Correlation between genetic distance and hAT (a), hAT.Ac (b) transposons, and the number of DNA (c) and hAT (d) transposon changes. The genetic distances were distance to outgroup Kurtiformes. Since all Gobioidei were derived from the outgroups at the same time, the genetic distances are served as a proxy for mutation rate. The correlation was conducted using Kendall’s ρ coefficient after correcting for phylogeny using phylogenetically independent contrasts (see text for details)

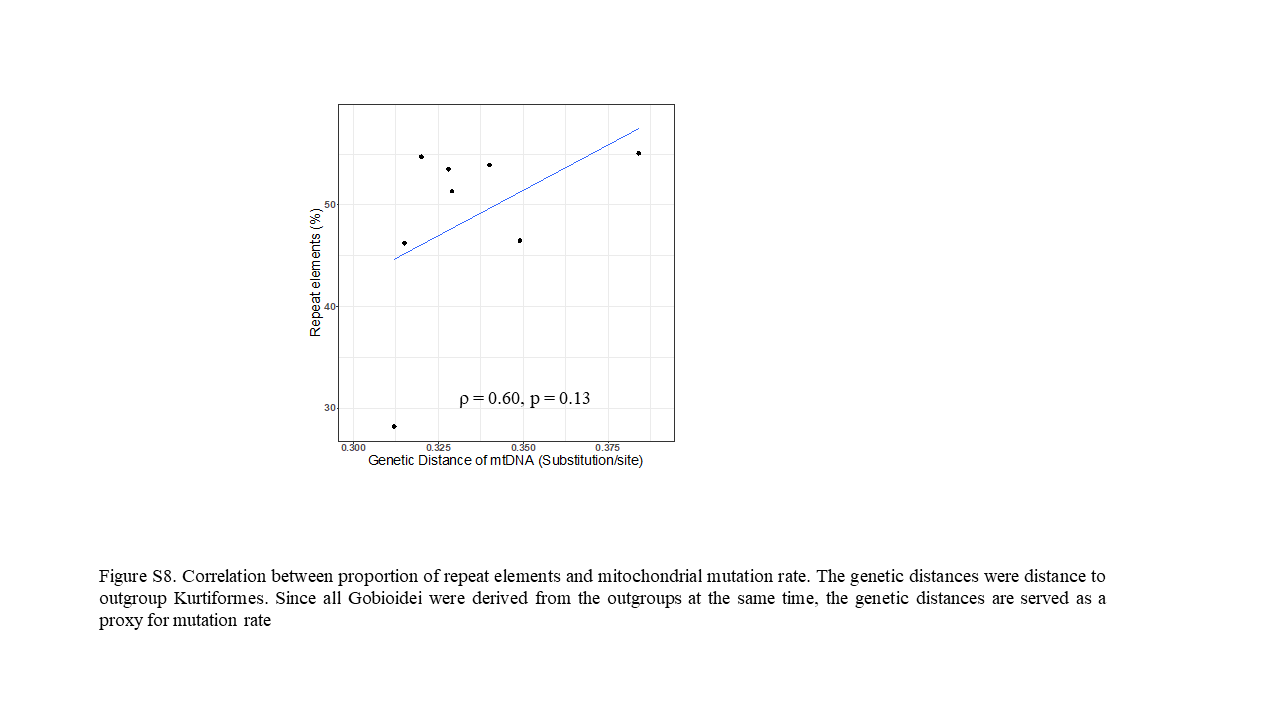

Figure S8. Correlation between proportion of repeat elements and mitochondrial mutation rate. The genetic distances were distance to outgroup Kurtiformes. Since all Gobioidei were derived from the outgroups at the same time, the genetic distances are served as a proxy for mutation rate
